# Comprehensive chromatin proteomics resolves functional phases of pluripotency

**DOI:** 10.1101/2022.08.08.503208

**Authors:** Enes Ugur, Alexandra de la Porte, Sebastian Bultmann, Micha Drukker, Matthias Mann, Michael Wierer, Heinrich Leonhardt

**Author notes:** These authors contributed equally (Alexandra de la Porte, Sebastian Bultmann). To whom correspondence should be addressed. Correspondence should be addressed to Heinrich Leonhardt, Michael Wierer or Matthias Mann.

## Abstract

The establishment of cellular identity is driven by transcriptional and epigenetic regulation exerted by the components of the chromatin proteome - the chromatome. However, chromatome composition and its dynamics in functional phases of pluripotency have not been comprehensively analyzed thus limiting our understanding of these processes. To address this problem, we developed an accurate mass spectrometry (MS)-based proteomic method called Chromatin Aggregation Capture (ChAC) followed by Data-Independent Acquisition (DIA) to analyze chromatome reorganizations during the transition from ground to formative and primed pluripotency states. This allowed us to generate a comprehensive atlas of proteomes, chromatomes, and chromatin affinities for the three pluripotency phases, revealing the specific binding and rearrangement of regulatory complexes. The technical advances, the comprehensive chromatome atlas, and the extensive analysis reported here provide a foundation for an in-depth understanding of mechanisms that govern the phased progression of pluripotency and changes of cellular identities in development and disease.

## Introduction

DNA- and chromatin-binding proteins regulate gene expression and thereby govern cellular identity. During early embryonic development, the chromatin of pluripotent stem cells (PSCs) undergoes dynamic changes that are conserved among mammals^1–5^. Pluripotency progresses in separate phases controlled by distinct signaling pathways and downstream transcription factors^3, 6, 7^. Three major intermediate phases of pluripotency have been described: naive (also referred to as ground state), formative, and primed^3^. Ground state PSCs harbor a homogeneously organized and transcriptionally permissive chromatin with high plasticity and low levels of repressive epigenetic marks^8, 9^. In transition to the formative phase, PSCs gain super-bivalency at promoters, and the exclusive ability to differentiate into primordial germ cells, while losing the expression of certain naive genes^1, 10^. Finally, at the primed phase, PSCs are partially fate determined, yet still share a core regulatory circuitry with earlier pluripotency phases^3, 11–14^.

Current systems-wide knowledge of pluripotency is primarily based on transcriptome and epigenome analyses, and chromatin accessibility data^1, 10, 14–16^. For instance, previous studies revealed that major chromatin reorganization and compaction occur at the formative phase^10^. However, how this chromatin reorganization affects chromatin proteome composition, the chromatome^17^, remains unknown. Moreover, although the expression of chromatin binders, such as transcription factors, has been extensively studied in PSCs^18–21^, changes in expression do not inevitably entail changes in chromatin association. Therefore, the complete picture of the chromatome structure and dynamics in functional phases of pluripotency is still largely missing.

Previous attempts to quantify global chromatomes combined high-resolution mass spectrometry (MS) with the biochemical purification of native^22, 23^ or formaldehyde (FA) crosslinked chromatin^24–27^. Although these methods greatly contributed to the understanding of the chromatome, they offer limited insights as they cannot detect low abundant DNA-binding factors that are known to play key regulatory roles despite low abundance. Furthermore, current sample preparation strategies require millions of cells (15-50 mio.) and multiple purification steps, which impairs overall protein recovery and quantification^25, 26^. Therefore, the current view of the chromatome remains incomplete.

To overcome these difficulties, we developed a method that combines a new streamlined chromatin purification strategy, Chromatin Aggregation Capture (ChAC), with Data-Independent Acquisition (DIA) MS-based proteomics, a powerful strategy for rapid, accurate, and reproducible proteomics analysis with a broad dynamic range that allows identification of low abundant proteins starting with 100-250K cells. Using this method we generated accurate and comprehensive chromatome maps of mouse naive, formative, and primed PSCs that cover 80% of transcribed chromatin binders in single MS runs. Our analysis of these datasets revealed striking chromatome changes between different functional phases of pluripotency and provided evidence for novel, low abundant chromatin binders that are dynamically regulated in pluripotency transitions. Additionally, by comparing the abundance of proteins in chromatomes and proteomes, we were able to infer chromatin reorganizations mediated by differential affinities or subcellular localizations. Finally, we applied this approach to chromatomes of human PSCs to provide a mouse-to-human comparison of the pluripotency chromatome. Collectively, we present a comprehensive atlas of proteomes and chromatomes for the three pluripotency phases, thus revealing previously unknown details about how cell identity governing proteins are recruited to or evicted from chromatin in the process of pluripotency transitions. We have made the datasets available and searchable on an interactive web application, accessible on: https://ugur-enes.shinyapps.io/Chromatome_Atlas/.

## Results

### Chromatin Aggregation Capture (ChAC) followed by data-independent MS acquisition (DIA) enables near-complete chromatome identification and high-precision quantification

We hypothesized that accurate and comprehensive chromatin proteomics could be accomplished by combining Chromatin Aggregation Capture (ChAC) with Data Independent Acquisition (DIA). The method comprises nuclei isolation and formaldehyde crosslinking followed by an initial chromatin enrichment step similar to the Chromatin enrichment for proteomics (ChEP) protocol^25^. This is followed by an additional purification based on the protein aggregation capture (PAC) technique^28^ to generate specific and pure chromatin fractions, and achieve highly accurate quantification by DIA-based MS using the DIA-NN software package^29^. Briefly, in DIA, all peptide precursors that fall into a predefined mass-to-charge (*m*/*z*) window are fragmented and acquired on the MS2-level compared to selecting the top N most abundant peptide ions in a typical Data-Dependent MS Acquisition experiment (DDA)^30–34^. The application of DIA is especially relevant for the analysis of enriched cellular structures that consist of highly repetitive structural elements such as nucleosomes. Here, DIA is much more sensitive and accurate for lower abundant proteins than the more semi-stochastic DDA-based approach^35, 36^. To improve chromatome quantification accuracy and comprehensiveness, we optimized the protocol, MS acquisition strategy (Supplementary Fig. 1a-c), and raw data analysis (Supplementary Fig. 1d-h) (Supplementary Table 1).

To benchmark the chromatome protocol, we performed ChAC-DIA in naive mouse embryonic stem cells (mESCs) and compared it to a recent ChEP-based chromatome data set of mESCs (PRIDE: PXD011782)^37^. ChAC-DIA identified over 2.5 times more proteins in half of the MS acquisition time (Fig. 1b). In addition, ChAC-DIA quantified proteins more reproducibly with median Coefficients of Variation (CVs) of 4% compared to 16% in the previous study (Fig. 1b and Supplementary Table 1). The CV differences were even more pronounced at the peptide ion level (Supplementary Fig. 1e).

**Fig. 1:**
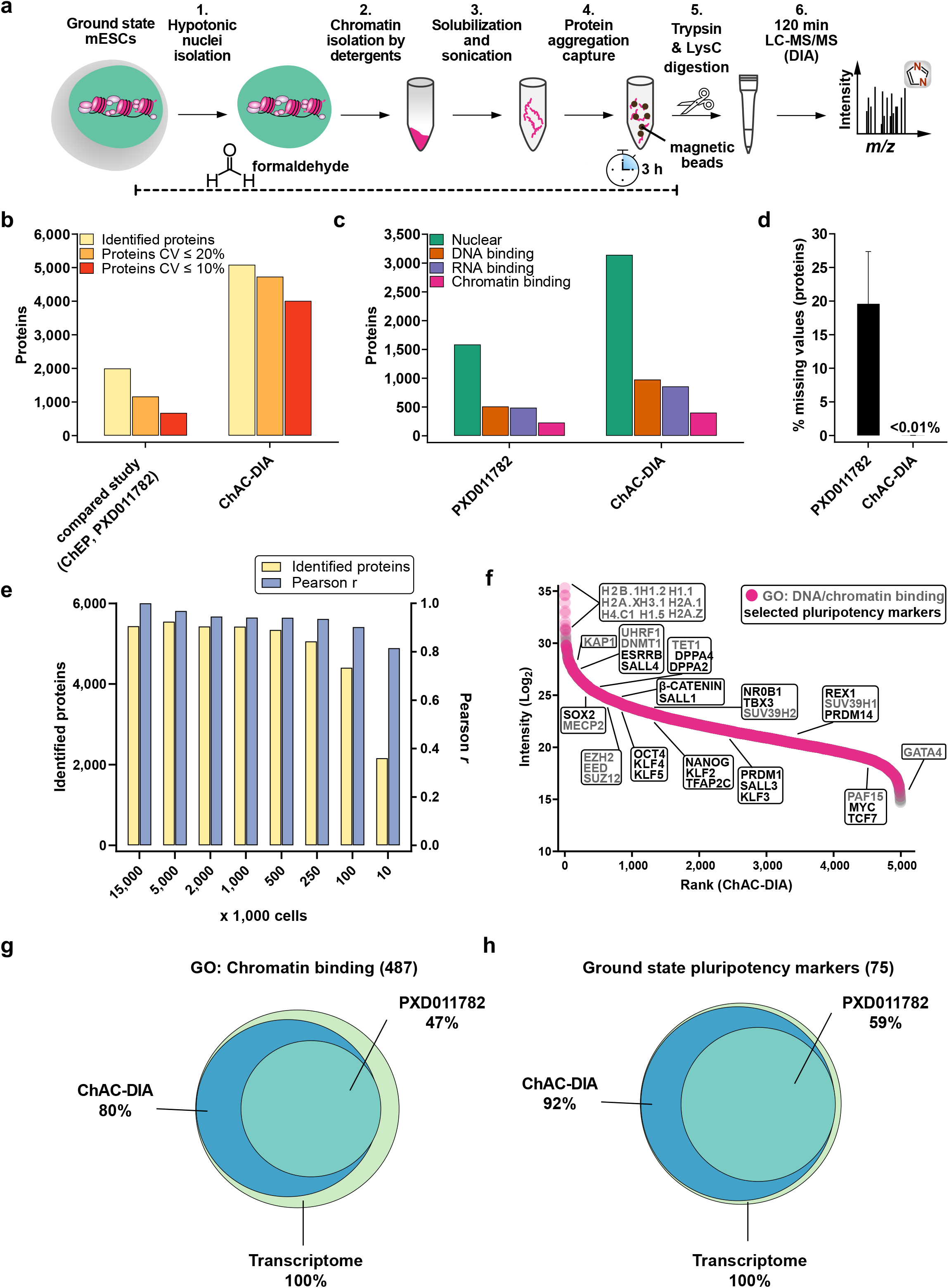
Chromatin Aggregation Capture (ChAC) followed by data-independent MS acquisition (DIA) enables near-complete chromatome identification and high-precision quantification. **a** Schematic workflow of ChAC-DIA. **b** Total numbers of identified proteins with representations of the coefficient of variation (CV) below 20% and 10%. ChAC-DIA results obtained in directDIA mode by DIA-NN were benchmarked against a previous study based on the ChEP protocol (PRIDE: PXD011782). In both cases, mouse naive PSCs were used. **c** Total numbers of proteins falling into a gene ontology (GO) category. **d** Percentage of missing intensity values on protein-level across replicates. **e** Total numbers of identified proteins and Pearson correlation coefficients of ChAC-DIA applied on different cell amounts. Pearson r reflects the correlation with the standard protocol comprising 15 Mio cells. **f** Protein abundance rank based on the ChAC-DIA-derived naive PSC chromatome. Chromatin binding proteins are highlighted in magenta. Protein names in black indicate examples of *bona fide* pluripotency factors. Protein names in gray indicate other chromatin binders and the highest ranked 9 proteins. **g** Venn diagram of proteins annotated as chromatin binding in ChAC-DIA, the compared study, and a transcriptome data set of naive PSCs (ArrayExpress: E-MTAB-6797). **h** Venn diagram of literature derived *bona fide* naive pluripotency factors identified by ChAC-DIA, the compared study, and a transcriptome data set of naive PSCs (ArrayExpress: E-MTAB-6797). See also **Supplementary Fig. 1,2**.

Next, we classified nuclear, DNA-binding, RNA-binding, or chromatin-binding proteins based on their Gene Ontology (GO) annotations^38^. ChAC-DIA identified more than twice the number of nuclear and DNA-binding proteins, and three times more unique peptides of DNA-binding proteins as the previous ChEP method despite half of the required MS time (Fig. 1c and Supplementary Fig. 1e). Furthermore, annotated chromatin proteins had significantly fewer missing values across replicates (Fig. 1d) and smaller CVs (Supplementary Fig. 1h).

To make the method applicable to rare stem cell populations, we examined how input amounts affect the performance of our method. Cell numbers between 100K to 5 Mio. correlated well with the original protocol comprising 15 Mio. cells (Pearson correlation > 0.9) and 250K to 5 Mio. cells were sufficient for stable identification rates of over 5,000 proteins (Fig. 1e). Notably, ChAC-DIA with as few as 10K cells still resulted in over 2,000 protein identifications. Ranking proteins quantified by ChAC-DIA according to their abundance revealed specific enrichment of histones and *bona fide* naive pluripotency factors as compared to a full proteome (Fig. 1f, Supplementary Fig. 2d, and Supplementary Table 1).

To further assess the comprehensiveness of ChAC-DIA, we compared the results to naive mESC transcriptome data. Among approximately 13,000 transcripts expressed in naive mESCs, 487 code for proteins annotated as chromatin binders, of which 80% were identified by ChAC-DIA (Fig. 1g). Among *bona fide* naive pluripotency factors, 92% were identified by ChAC-DIA. Given that not all transcripts are translated into proteins with the same efficiency, we also compared the results obtained by ChAC-DIA to a full proteome analysis covering around 7,000 proteins, and observed that ChAC-DIA identified the same number of known chromatin binders that were also present in the full proteome data (Supplementary Fig. 2a-c). Taken together, our results validated ChAC-DIA as a rapid and highly accurate method for analyzing the chromatome that uses only 100-250K cells and achieves unprecedented, almost complete chromatome coverage, including low abundant proteins.

### Chromatome mapping reveals a specific enrichment of chromatin-associated proteins in ground state PSCs

To define high confidence chromatomes of ground state PSCs and thereby assess the specificity of chromatin enrichment by ChAC-DIA, we analyzed all fractions obtained during the chromatin purification in triplicates (i.e. whole cell lysate, cytoplasmic and nuclei fractions, ChAC-DIA after 1-3 washes). In total, we identified 8,567 proteins, and the triplicates correlated well with each other (R^2^ > 0.95). We observed that the correlation between the chromatin and nuclei fractions was weak (R^2^ = 0.66) (Supplementary Fig. 3a-d). Filtering for proteins with significantly different quantities between the fractions (ANOVA FDR < 0.05, fold change difference ≥ 1.5), resulted in 5,464 proteins which explains the low correlation between the fractions. Unsupervised hierarchical cluster analysis of these proteins revealed nine distinct clusters (Fig. 2a and Supplementary Table 2).

**Fig. 2:**
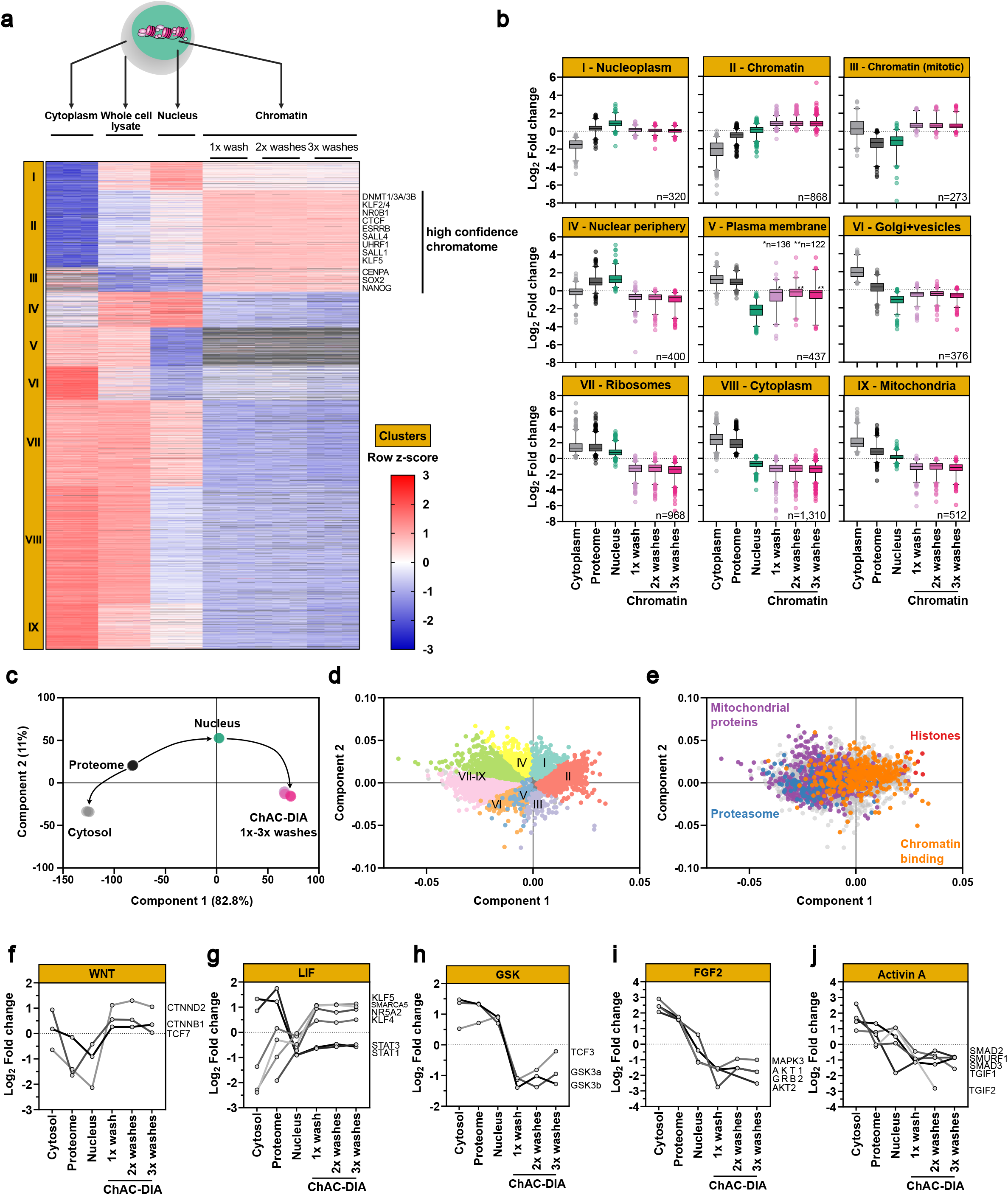
Chromatome mapping reveals a specific enrichment of chromatin-associated proteins in ground state PSCs. **a** Different fractions along the ChAC-DIA protocol were processed and measured. After ANOVA testing (FDR < 0.05, fold change difference ≥ 2) results were visualized in a heatmap generated by unsupervised hierarchical k-means clustering of z-scored intensities. In total 9 clusters were identified. Proteins that are enriched only in the chromatome fractions are highlighted as the high confidence chromatome. **b** Boxplot representation of row-scaled fold changes within each cluster. Cluster names are based on the most prominent GO enriched terms (see Supplementary Fig. 3 e). **c** Principal component analysis (PCA) of the six different fractions. **d, e** Loadings of the PCA. Proteins are colored according to clusters (**d**) or GO categories (**e**). **f-j** Individual intensity profile plots of several proteins that are components of the WNT, LIF, GSK, FGF2, or Activin A pathways. See also **Supplementary Fig. 3**.

Two clusters (II and III), harboring 1,141 proteins, were significantly enriched in the chromatomes (ChAC-DIA after 1-3 washes), but not in the nuclei or any other fraction. Therefore, proteins in clusters II and III comprise high-confidence chromatin binders. Importantly, well-known pluripotency proteins such as DNMT1, ESRRB, SALL4, or SOX2 are most abundant within these two clusters. Cluster II contained the highest enrichment of general chromatin-specific GO categories such as “nucleosome” or “nucleosomal DNA binding” (Supplementary Fig. 3e and Supplementary Table 2). Euchromatic and heterochromatic proteins were equally enriched within this cluster. In cluster III, mitotic chromatin binders were overrepresented, resulting in GO categories such as “mitotic prometaphase”. Clusters I and IV revealed significant enrichment of proteins in the nuclei fraction and a strong depletion in the chromatomes indicating that these two clusters captured nucleoplasmic proteins (Fig. 2b). In line with this, well-characterized nucleoplasmic proteins such as RANGAP1 or CDK11B were categorized within these two clusters. In contrast, proteins in clusters V-IX were enriched for cytoplasm-specific GO categories (e.g. “Golgi membrane” or “structural constituent of the ribosome”). Individual loadings of the PCA (Fig. 2c) uncovered a distinct separation for each identified cluster (Fig. 2d) and, thus, confirmed that chromatin-related proteins were mostly enriched in clusters II and III whereas clusters V-IX harbored most mitochondrial or proteasome-related proteins (Fig. 2e and Supplementary Fig. 3f).

Pluripotency phases are governed by distinct signaling pathways that lead to the translocation of otherwise cytoplasmic transcription factors into the nucleus^39–42^. For example, naive pluripotent stem cells harbor active WNT and LIF pathways, while the GSK, FGF2, and Activin A pathways are inactive. Our data captured these features accurately, as we observed the chromatin-association of transcription factors linked to the WNT and LIF pathways, while those related to GSK, FGF2, and Activin A were mostly cytoplasmic (Fig. 2f-j). For instance, β-CATENIN, the effector of WNT signaling, was equally distributed between the cytoplasmic and chromatin fractions, while being less abundant in the nuclear fraction (Fig. 2f). We also observed chromatin enrichment of the LIF pathway transcription factors like KLF4 and KLF5, as well as STAT1 and STAT3, which, although being less abundant at chromatin than in the cytoplasm, still showed chromatin enrichment over the nuclear fraction (Fig. 2g). In contrast, GSK, FGF2, and Activin A related transcription factors were depleted from the chromatin fractions (Fig. 2h-j). Taken together, we confirmed that ChAC-DIA selectively enriched components of the chromatome by reducing background proteins, even hard to separate mitochondrial or ribosomal proteins. This enabled identification of not only the majority of the annotated chromatome, but expansion of the existent GO annotations. Thus, ChAC-DIA provides a high-confidence global map of the chromatome. Furthermore, analyzing chromatome data in combination with the overall proteome, and proteomes derived from different cellular fractions, allowed us to dissect events such as nuclear translocation and chromatin binding of proteins related to pluripotency-regulating pathways.

### Chromatome atlas of mouse naive, formative, and primed pluripotent stem cells identifies groups of chromatin proteins with distinct binding patterns

Two recent studies provided evidence that the formative phase is a discrete pluripotent state during embryonic development that is transcriptionally distinct from naive and primed pluripotency phases^1, 10^. To examine this further, we analyzed chromatomes of naive, formative, and primed PSCs (Fig. 3a). We observed that 1,403 proteins significantly changed in the chromatome during the differentiation of naive to formative PSCs, while the proteome revealed 1,683 significantly regulated proteins (*p* value < 0.05, FC ≥ 2) (Fig. 3a). In contrast, between formative and primed PSCs, only 859 proteins were significantly regulated on chromatome level and 1,451 on proteome level. This suggests a more drastic reorganization of the chromatome during the transition from naive to formative pluripotency.

**Fig. 3:**
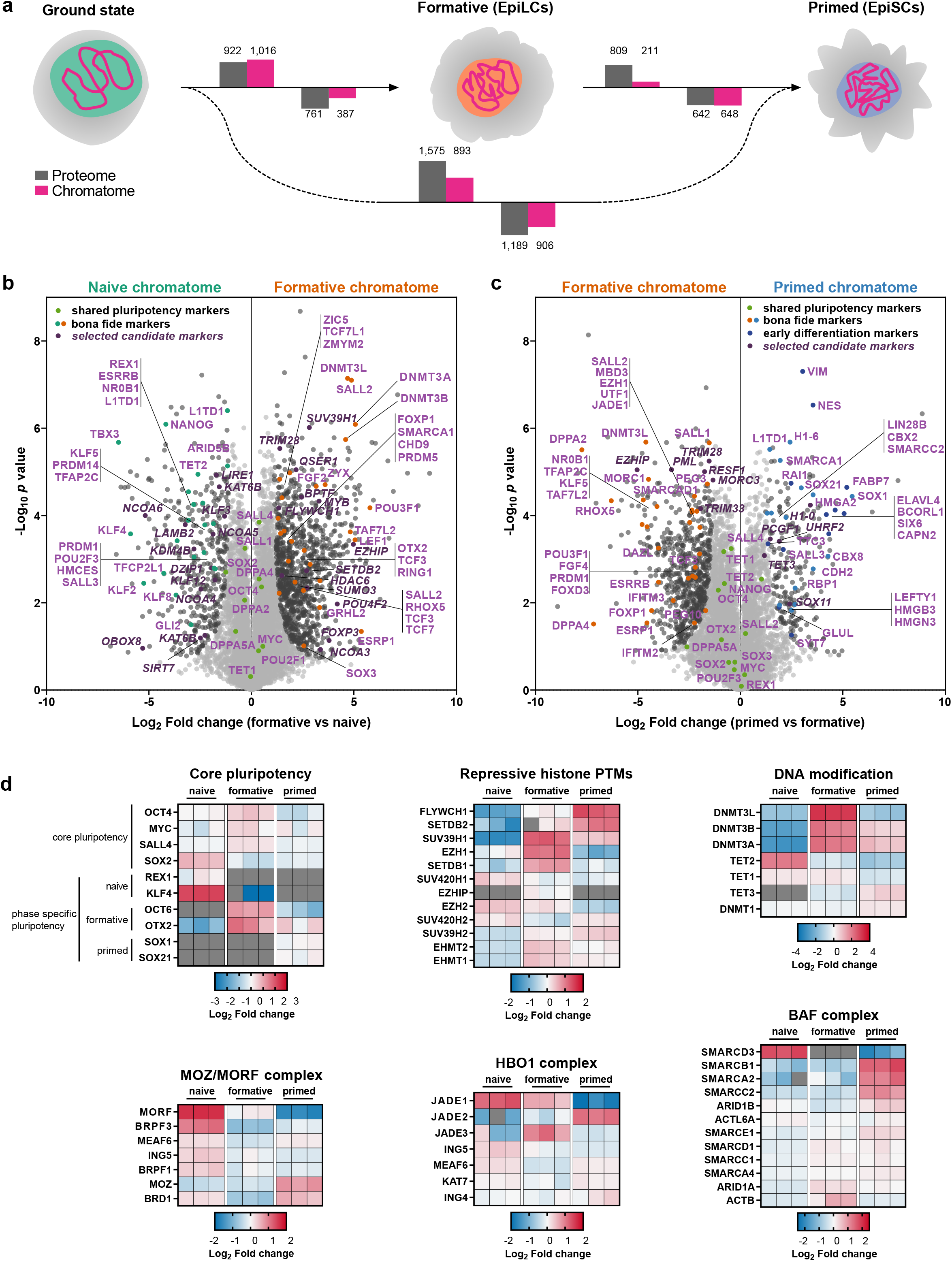
Chromatome atlas of mouse naive, formative, and primed pluripotent stem cells identifies groups of chromatin proteins with distinct binding patterns. **a** Schematic representation of compared cell lines and total significant changes between respective proteomes and chromatomes (Student’s T-test, *p* value < 0.05, FC ≥ 2). **b, c** Volcano plots of chromatomes based on Student’s T-test displayed in (**a**). Light grey dots: not significantly enriched proteins. Black dots: significantly enriched proteins. Green dots: shared pluripotency factors. Blue dots: early differentiation markers. n = 3 biological replicates, meaning independently cultured/differentiated PSCs of the same genetic background. **d** Heatmap representation of pluripotency factors and proteins related to repressive histone PTMs, DNA modifications as well as constituents of the MOZ/MORF, HBO1, and BAF complexes. Fold changes are row-normalized by subtracting the mean log_2_ fold-change from each value. See also **Supplementary Fig. 4-9**.

Next, we analyzed the chromatome changes based on a list of PSC phase-specific factors that we derived from the literature (Supplementary Table 3)^1, 4, 6, 7, 10, 13, 15, 37, 43–51^. ChAC-DIA data confirmed that the abundance of the core pluripotency circuitry (OCT4, MYC, SOX2, and SALL4) is maintained throughout pluripotency; whereas state-specific markers displayed phase-dependent selective enrichment in the chromatome (Fig. 3b-d and Supplementary Table 3). The naive chromatome was characterized by high levels of REX1, ESRRB, KLF4, and TET2 while the *de novo* methyltransferases DNMT3A and DNMT3B, OTX2, and OCT6 (or POU3F1), were highly enriched in the formative chromatome. Notably, most differences either occurred between formative and naive, or formative and primed PSCs. However, several proteins that appeared significantly increased during naive to formative transition were strikingly decreased when comparing formative vs primed PSCs, suggesting a formative-specific function for these proteins. Besides classical formative pluripotency factors such as OCT6^13, 52^ this group of proteins also contained DNMT3L, which was recently postulated to be a putative marker of formative pluripotency^43^, as well as proteins without a reported pluripotency association, like QSER1, EZHIP, and SUV39H1. In addition, we observed a slight enrichment of lineage-specific transcription factors such as NES as early as the formative state.

In contrast to the formative chromatome, the primed chromatome was characterized by lower levels of early post-implantation-specific proteins like DPPA4^15^ and OCT6^7^ and higher levels of *bona fide* primed-specific transcription factors such as SOX1^10^ and SALL3^43^. Similarly, naive factors like ESRRB, HMCES, and TET2 were decreased in the primed chromatome while lineage-specific factors such as RAI1 and SIX6 (Fig. 3c) were significantly enriched, which fits the partially fate-determined identity of primed PSCs. Among the primed-specific chromatin constituents, several histone H1 variants and high mobility group (HMG) proteins were also observed. The enrichment of these proteins governing chromatin structure and compaction could in part account for the previously described reduced chromatin plasticity and accessibility at the primed phase^1, 5, 10^. Although major chromatome changes were already established at the formative state, these results demonstrate that formative and primed pluripotency are characterized by distinct chromatin landscapes.

Of note, even very low abundant chromatin binders like FLYWCH1, which were not detected in previous chromatome or proteome studies of PSCs, were readily detected and quantified by ChAC-DIA^43, 53^. FLYWCH1 was recently described to bind H3K9me3-rich regions in HeLa cells^53^. Interestingly, we observed that FLYWCH1 successively increases at chromatin from naive to primed PSCs when also H3K9me3 increases as determined by PTM analysis on the ChAC-DIA data (Supplementary Fig. 6c).

These findings point to gradual chromatin recruitment or eviction of pluripotency governing factors during naive to primed transition. Interestingly, we observed similar chromatin-enrichment patterns for proteins related to epigenetic regulation, transcriptional regulation, and chromatin remodeling, as well as hundreds of zinc finger proteins with mostly unknown functions in pluripotency regulation (Fig. 3d). Approximately 70% of proteins harboring a zinc finger domain significantly change between naive and primed pluripotency, which fits well with the recently reported zinc finger protein-driven regulation of transposable elements during early embryonic development^54, 55^. In summary, we provide the first systematic and near-comprehensive chromatome data set of naive, formative, and primed PSCs as an atlas of pluripotency (Supplementary Fig. 4-8 and Supplementary Table 3). We show that the chromatome reflects distinct features of pluripotency phases and a tightly regulated pluripotency phase transition process.

### Determination of relative chromatin binding reveals regulatory changes along pluripotency phases

Next, we correlated the transcriptome changes during the naive to formative transition^56^ with the respective proteome and chromatome changes. As expected and previously reported^43, 57, 58^ the proteome showed a moderately positive correlation with the transcriptome (Fig. 4a), due to mechanisms regulating translation and protein stability. Interestingly, a similar difference was found in the comparison of proteome and chromatome changes (Fig. 4b), pointing to mechanisms controlling chromatin binding and dissociation. Consequently, transcriptome and chromatome showed the lowest correlation (Fig. 4c) indicating that transcriptional data can only provide limited coverage of regulatory chromatin changes. In line with these observations, proteins related to active signaling pathways in postimplantation pluripotency like the FGF2, Activin A, and Notch pathways were differentially enriched in the chromatome, while they changed neither on transcriptome nor on proteome level.

**Fig. 4:**
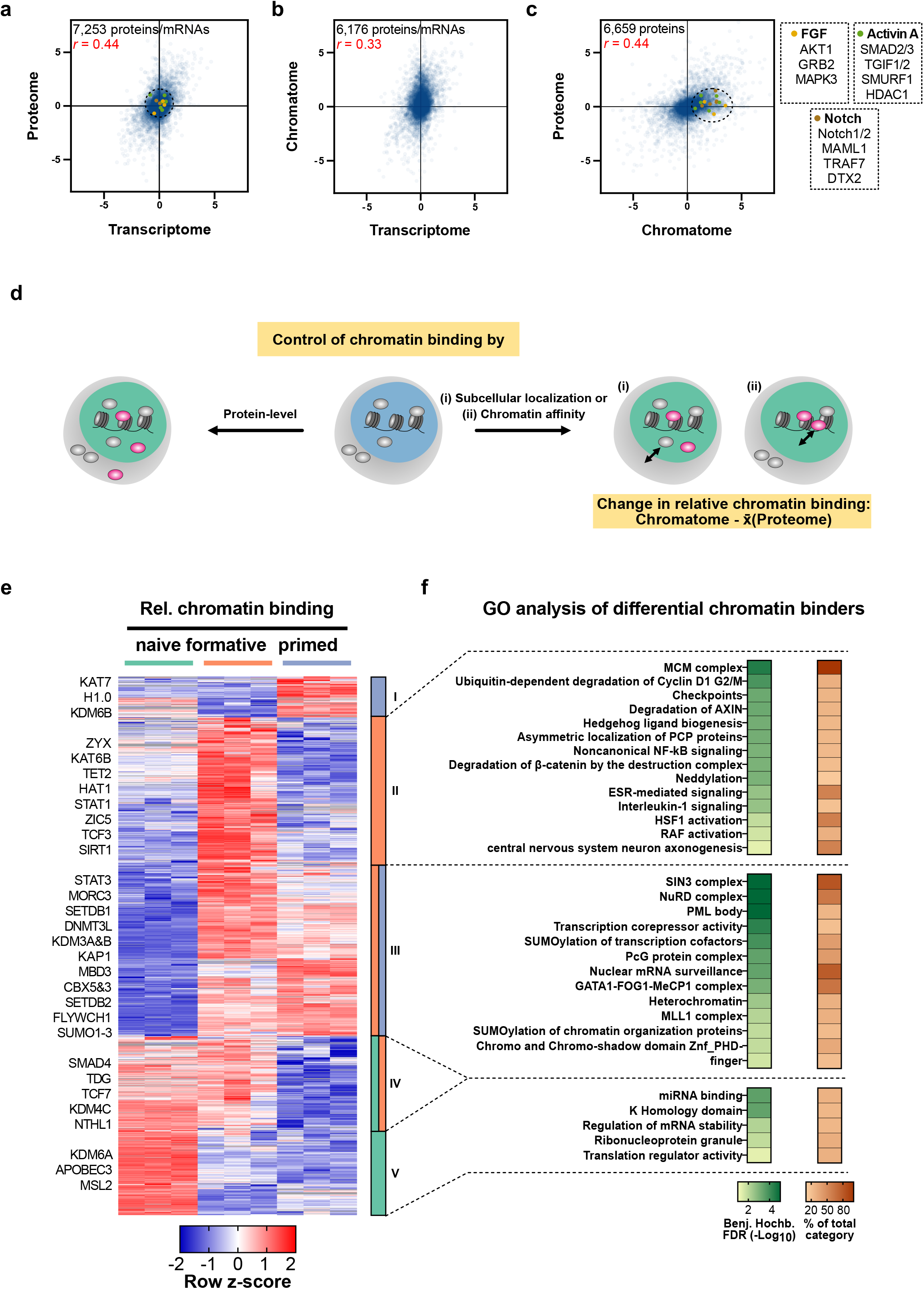
Relative chromatin binding reveals higher chromatin affinity of heterochromatic proteins in formative and primed PSCs. **a-c** Correlations of transcriptomes (ArrayExpress: E-MTAB-6797), proteomes, and chromatomes of formative vs naive PSCs of isogenic background (J1). Only proteins/mRNAs that were identified in both compared data sets are displayed. Pearson correlation coefficients are indicated in red. **d** Schematic representation of the relative chromatin binding concept. **e** Row z-scored relative chromatin binding changes between naive, formative and primed PSCs filtered for ANOVA significant changes (FDR < 0.05, FC ≥ 2) and high-confidence chromatin binders. The relative chromatin binding was computed by subtracting the FC on chromatome level by the mean proteome FC of either formative vs naive or primed vs formative PSCs, respectively. **f** GO analyses of proteins enriched in cluster II, III or V of the hierarchical clustering from (**e**). As a comparison the whole set of identified proteins were utilized.

Proteome-independent changes in the chromatome contain valuable information, and point to either altered chromatin affinity or subcellular localization and availability of individual proteins (Fig. 4d). We therefore computed proteome normalized chromatome changes to estimate the relative changes in chromatin binding. We subtracted the Log_2_ chromatome-intensity of a protein from its mean Log_2_ proteome intensity across triplicates and subsequently filtered for significant proteins by ANOVA testing (FDR < 0.05 and FC > 2) (Fig. 4e and Supplementary Table 3). Based on our differential chromatin fraction analysis, we defined high confidence chromatin binders as proteins that are significantly enriched in the chromatome over the proteome.

We observed that 1,518 proteins significantly changed in relative chromatin binding from naive to primed pluripotency. Hierarchical clustering yielded five distinct clusters harboring proteins with different trends in relative chromatin binding across pluripotency phases. GO analysis of these five clusters against the background of total identified proteins revealed distinct functional categories (Benjamini-Hochberg FDR < 0.05) (Fig. 4f and Supplementary Table 3). In the cluster of proteins with a peak in relative chromatin binding at the formative phase (cluster II) categories related to signaling pathways like “ -catenin degradation” or “RAF activation” were enriched (Fig. 4f). Importantly, cluster III showed an increased relative chromatin binding at the formative and primed phases and was enriched for categories associated with a repressive chromatin state like “heterochromatin” or “transcription corepressor activity”. More specifically, this cluster harbored essential heterochromatic proteins such as SETDB1, SETDB2, KAP1, CBX3, and CBX5 suggesting a functional relation of their formative and primed specific enrichment to the incremental heterochromatinization towards the exit from pluripotency. Interestingly, this cluster III was also enriched for GO categories related to “SUMOylation of transcription factors”, “SUMOylation of chromatin organization proteins”, and SUMOylation dependent “PML bodies”. In line with this observation, SUMOylation was reported to regulate heterochromatinization in naive mouse PSCs^59^. Notably, histone H1.0, whose function in chromatin compaction depends also on its SUMOylation^60^, peaked in its relative chromatin binding at the primed phase. These results suggest that besides the binding of classical heterochromatin factors, SUMOylation also contributes to heterochromatin formation at the formative and primed phases. Among the proteins with decreasing relative chromatin binding (clusters IV and V) are enzymes involved in DNA and histone demethylation like TDG, APOBEC3, NTHL1, KDM4C, and KDM6A. Thus, lower levels of these proteins would translate into an increase of repressive epigenetic marks, which is expected to promote repressive chromatin states and reduce chromatin plasticity.

These findings show that a combined analysis of proteomes and chromatomes may reveal changes in chromatin affinity along with cellular differentiation processes and thus pinpoint specific regulatory events.

### The chromatome of conventionally cultured human ESCs is most similar to the mouse primed state

Previous reports provided strong evidence that the epigenome, transcriptome, and proteome of conventional human ESCs (hESCs) correspond to post-implantation mouse PSCs^43, 61–63^. Here, we used our method to examine the correspondence between different pluripotency states of hESCs and mouse PSCs. A Venn diagram representation of the high confidence chromatomes for all three mouse PSCs and hESCs revealed an overlap of approximately 75% (Fig. 5a and Supplementary Table 4). The strongest overlap was between proteins related to chromatin remodeling, histone modifications, and developmental processes (Supplementary Fig. 9a). A PCA of the high confidence chromatomes resulted in a clear separation of all three mouse PSCs from hESCs on PC1. hESCs and mouse formative and primed PSCs in turn grouped further apart from mouse naive PSCs on PC2 (Fig. 5b, see also Supplementary Fig. 9b). Interestingly, the high confidence chromates of the human and mouse post-implantation stages grouped even more closely at the level of relative chromatin binding (Fig. 5c).

**Fig. 5:**
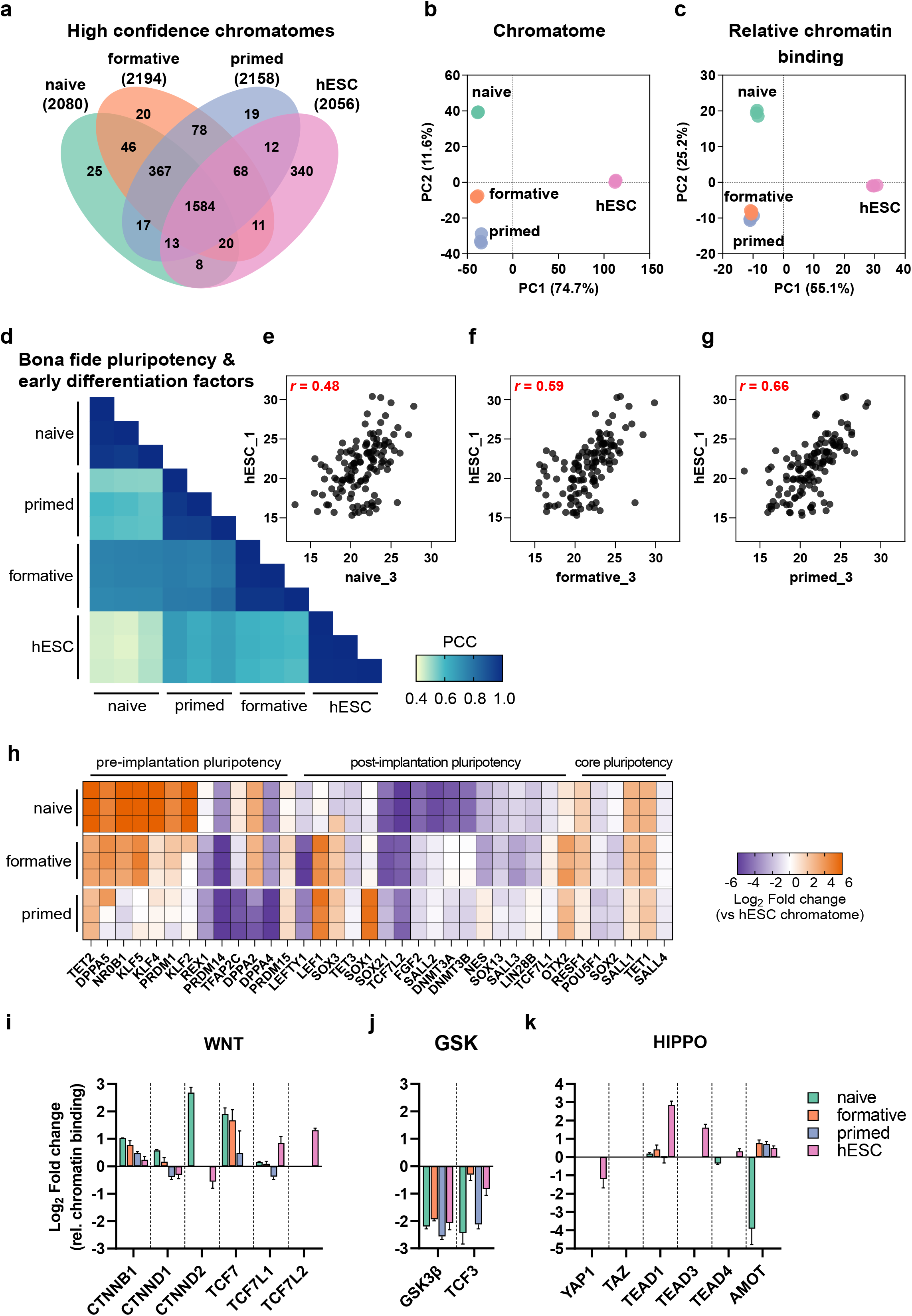
The chromatome of conventionally cultured human ESCs is most similar to the mouse primed state. **a** Venn diagram of high confidence chromatomes in all tested cell lines. The high confidence chromatome was defined by a Student’s T-test between each cell line’s chromatome vs proteome (Student’s T-test, *p* value < 0.05 and FC ≥ 1.5). **b, c** PCA of the high confidence chromatomes of the tested four cell lines on chromatome (**b**) and relative chromatin binding-level (**c**). **d** Pearson correlations of chromatomes filtered for literature-derived *bona fide* pluripotency and differentiation factors. **e-g** Scatter plots of one replicate of hESCs vs naive (**e**), formative (**f**) or primed PSCs (**g**) and Pearson correlation coefficient from (**d**) are displayed in red. **h** hESC-normalized chromatomes from each mouse PSC to hESCs in Log_2_. The selection comprises *bona fide* pluripotency factors. **i-k** Relative chromatin bindings of proteins related to the WNT, GSK or HIPPO signaling pathways in all analyzed cell lines. See also **Supplementary Fig. 9**.

To further dissect whether hESCs correspond more to the early or late mouse post-implantation stage, we computed correlations between the chromatomes of all four cell lines selected for *bona fide* pluripotency and early differentiation factors (Fig. 5d). We noted an incremental increase in the correlation of hESCs with naive, formative, and primed PSCs (Pearson, *r* = 0.48 for naive, 0.59 for formative, and 0.66 for primed PSCs) (Fig. 5e-g), while chromatomes of formative and primed PSCs correlated better to each other (Pearson, *r* = 0.78) than to naive PSCs (Pearson, *r* = 0.74 and *r* = 0.57, respectively). We observed similar differences on the relative chromatin binding and total proteome levels (Supplementary Fig. 9c, d).

For an in-depth view of pluripotency factors and their contribution to cell identity, we computed the chromatome difference between a given mouse PSC-line and hESCs for each *bona fide* pluripotency factor (Fig. 5h and Supplementary Table 4). A step-wise loss of pre-implantation pluripotency markers was observed from naive to primed PSCs with some remarkable exceptions; TFAP2C, DPPA2, DPPA4, and PRDM14 were more similar in their chromatin abundance between both naive and formative PSCs and hESCs. These proteins are indicative of germline competence, a capability that only mouse formative PSCs and conventional hESCs harbor^51, 64–67^. Moreover, REX1, a well-characterized naive pluripotency and germline marker, was more strongly associated with the hESC chromatome than mouse formative and primed PSCs, likely reflecting the more heterogeneous nature of hESCs or species-specific differences^68^. In a PCA based on these *bona fide* pluripotency factors only, mouse formative PSCs were even further separated from primed PSCs but not from naive PSCs (Supplementary Fig. 9e, g). The loadings of the PCA uncovered that the main causes of this strong separation were naive pluripotency factors such as NR0B1, KLF2, and KLF4 (Supplementary Fig. 9f). Thus, these naive factors were less associated with chromatin in hESCs and mouse primed PSCs than formative or naive PSCs. Conversely, post-implantation pluripotency factors contributed to the higher similarity between hESCs and primed PSCs. Of note, we did not observe differences in the chromatin association of the core pluripotency circuitry such as OCT4 or SALL4 (Supplementary Fig. 9g, h).

Contrasting the relative chromatin binding of pluripotency governing signaling pathways also revealed a higher similarity of hESCs and mouse formative and primed PSCs considering e.g. the WNT and GSK pathways (Fig. 5i, j). However, there were some striking differences in the HIPPO pathway (Fig. 5k). This pathway is highly active in pluripotent epiblast cells and upon its activation the downstream proteins YAP1 and TAZ are kept cytoplasmic^39, 69^. Interestingly, we observed YAP1 and TAZ only in the full proteome fractions, except for hESCs where YAP1 was also present in the chromatin fraction. This was in agreement with a higher relative chromatin binding of the YAP1 cofactors TEAD1/3/4 in hESCs, likely suggesting a more inactive state of the HIPPO pathway in hESCs than in closely related mouse pluripotency phases.

In summary, by additional analysis of hESCs, we have shown that the proteome, chromatome, and relative chromatin binding in hESCs are reminiscent of mouse post-implantation-reflecting PSCs and mouse primed PSCs, with differences that could be attributed to variances between the species or between the actual pluripotency phases.

## Discussion

Previous studies have established methods for chromatin purification and measurement^24–27, 70, 71^. These techniques, however, require large numbers of cells and have limited accuracy and comprehensiveness, often failing to detect low abundant proteins such as regulatory factors. In this study, we combined a stringent and simple chromatin preparation strategy of crosslinked nuclei with an additional purification step by protein aggregation capture (PAC) and optimized DIA-based MS. Requiring only three hours of experimental hands-on time, our method confidently reduces non-chromatin proteins while identifying more than twice the number of DNA-binding proteins compared to other methods in half of the MS acquisition time^37, 72^. In addition, recent deep neural network-based computational processing of DIA measurements without a peptide library (direct DIA) can now outperform DDA in accuracy and comprehensiveness^29, 33, 73, 74^. Thus, our direct DIA measurements additionally decreased instrument time, while providing a near-complete chromatome coverage. However, it is possible that a library-based analysis would increase the current chromatome depth further, and may represent a potential future opportunity.

The datasets generated here allowed us to perform several different types of analysis. Given that ChAC-DIA selectively enriched components of the chromatome, we were able to assemble a high-confidence global map of the chromatome. By comparing chromatome and proteome data, including proteomic data derived from different cellular fractions, for different pluripotency phases, we identified proteins affected by nuclear translocation or chromatin binding. For example, we observed chromatin enrichment of cytoplasmic transcription factors such as those involved in WNT and LIF pathways, and not GSK, FGF2, and Activin A pathways in naive PSCs, which has implications for their role in pluripotency regulation. Furthermore, normalizing the chromatome to protein levels enabled a global assessment of changes in relative chromatin binding which may be caused by either altered chromatin affinity and accessibility or differential subcellular localization and availability. Our method thus enables accurate and comprehensive chromatome and relative chromatin binding measurements despite limited cell numbers, making it ideally suited for analyzing minute tissue samples or rare subpopulations of cells.

Additionally, ChAC-DIA enables quantification of low-abundant transcriptional or epigenetic regulators, and we identified several low-abundant chromatin binders that are pluripotency phase-specific. Besides well-described factors, we find many phase-specific proteins with still unknown functions in pluripotency regulation. Given their phase-specific chromatin association, many of them are likely to contribute to the regulation of cellular identity. One such example is EZHIP which was only identified in the formative phase. EZHIP was recently described to inhibit H3K27me3 by mimicking the H3K27M oncohistone and thus preventing the PRC2 complex from spreading along chromatin^75, 76^. Bulk levels of H3K27me3 are known to be downregulated from naive to primed pluripotency while super-bivalent sites harboring H3K4me3 and H3K27me3 are enriched^10, 77^. In our chromatome data set, we observed that EZH1 increases at chromatin between the naive and formative PSCs which does not fit a global downregulation of H3K27me3. Interestingly, this goes along with an increase in EZHIP in the formative chromatome implying a possible role of PRC2 inhibition or redirection to other regions by EZHIP in formative PSCs. Moreover, low abundant epigenetic writers such as SUV39H1/2, SUV420H1/2, SETDB2, or TET1-TET3 featured phase-specific enrichment at chromatin. Remarkably, all three TET proteins showed a distinct redistribution along the exit from pluripotency, starting with TET1 and TET2 being most abundant in the naive state and TET3 being mostly chromatin-associated in the primed state. This was also observed in conventional hESCs where TET2 and TET1 are even less associated with chromatin than in mouse primed PSCs.

The chromatome correlates weakly with the transcriptome and proteome and is, therefore, an important complement to previous studies of pluripotency. Our results provide a system-wide view of pluripotency by offering a chromatome atlas with specifically enriched proteins for each analyzed pluripotency phase. Our observations are in line with the recent finding that formative pluripotency is an essential state which is transcriptionally and epigenetically distinct from naive pluripotency and to a smaller degree also from primed pluripotency^1, 3, 10, 13, 45, 78^. The underlying chromatome changes fit in with the phased progression model of pluripotency^3^. Moreover, formative and primed PSCs share the majority of open chromatin sites while there is little overlap between formative and naive PSCs^1^. Our data support this observation by showing that the chromatome undergoes larger changes from naive to formative, than from formative to primed pluripotency. The chromatin composition is further reorganized between formative and primed PSCs, mainly driven by transcription factors triggering early differentiation as well as histone H1 and HMG variants guiding chromatin compaction. The histone H1 chromatin enrichment is in agreement with an increased relative chromatin binding of SUMO1-3 and SUMOylating enzymes of chromatin organizing proteins. SUMOylation of histone H1 was recently described as a mechanism for heterochromatinization in ESCs^60^, thus suggesting a role for SUMOylation in further chromatin compaction from formative to primed pluripotency. An increased relative chromatin binding was observed for additional heterochromatic proteins, such as KAP1 and CBX3, at the formative and primed phases. We conclude that heterochromatic proteins not only change in abundance at later pluripotency phases but also display a higher chromatin affinity that fits in with increased levels of repressive epigenetic marks.

Conventionally cultured hESCs are reminiscent of mouse primed PSCs regarding their epigenome, transcriptome, and underlying signaling cues^39, 61^. Still, human embryonic development comprises pluripotent phases that differ in length and growth conditions when compared to mouse^1, 3, 4, 79–81^. It remains unclear whether hESCs are the direct counterpart of mouse primed PSCs and to what extent they share unique features with mouse formative PSCs. A quantitative comparison of the high confidence chromatomes revealed that mouse primed PSCs correlated best to hESCs. Of note, a comparable correlation range was previously described on transcriptome and full proteome levels^43, 51^. In our hands, the correlation between hESCs and mouse primed PSCs increased even further when only *bona fide* pluripotency and early differentiation factors were considered. Here, chromatome-levels of naive pluripotency factors were the main difference between mouse primed PSCs and hESCs on the one side and mouse formative and naive PSCs on the other side. One major distinction between hESCs and mouse primed PSCs was the high chromatin association of essential germline factors like DPPA2, PRDM14, and TFAP2C in hESCs which resembles formative pluripotency in the mouse. This finding may explain the differential developmental capacities of hESCs and mouse primed PSCs. In addition, the hESC chromatome provided evidence for a less active HIPPO pathway compared to all three mouse PSCs, likely reflecting more species-specific signaling mechanisms.

Our study sheds light on the important question of whether cell identity-defining transcription factors coexist, suggesting an ongoing competition with each other^82, 83^, or abruptly change across pluripotency phases^4^. For all three phases and especially for the formative phase we observed that transcription factors were gradually recruited or evicted from chromatin. For instance, OTX2, a key transcription factor of formative pluripotency^15, 84^, peaks in abundance at the formative state, but is still associated with chromatin in naive and primed PSCs. Thus, our findings support the model of coexisting phase-specific transcription factors that ultimately define cellular identity if a certain critical threshold is exceeded.

In conclusion, we present a robust chromatin proteomics method to detect changes in abundance and affinity of even low abundant proteins. We offer a rich resource for the proteomes, chromatomes, and relative chromatin bindings in mouse naive, formative, and primed PSCs, as well as hESCs that are a basis for identifying and investigating novel regulatory mechanisms of pluripotency. Further investigations of candidate phase-specific proteins highlighted herein may help detangle the connection between pluripotency and lineage priming with an expected relevance for clinical applications of iPSCs. The dramatically improved sensitivity now makes it possible to also study rare subpopulations of cells. The comprehensive capture of chromatomes and chromatin affinities provides a deep and unbiased view into regulatory events underlying the establishment, maintenance, and change of cellular identity.

## Methods

### Cell culture

Naive J1 mESCs were cultured in serum-free media consisting of: N2B27 (50% neurobasal medium (Life Technologies), 50% DMEM/F12 (Life Technologies)), 2i (1 μM PD032591 and 3 μM CHIR99021 (Axon Medchem, Netherlands)), 1,000 U/mL recombinant leukemia inhibitory factor (LIF, Millipore), and 0.3% BSA (Gibco), 2 mM L-glutamine (Life Technologies), 0.1 mM β-mercaptoethanol (Life Technologies), N2 supplement (Life Technologies), B27 serum-free supplement (Life Technologies), and 100 U/mL penicillin, 100 μg/mL streptomycin (Sigma). Formative EpiLCs were derived by differentiating naive mESCs^51^ for 48 h using the same serum-free media for naive mESCs devoid of 2i, LIF, and BSA and supplemented with 10 ng/mL Fgf2 (R&D Systems), 20 ng/mL Activin A (R&D Systems) and 0.1 × Knockout Serum Replacement (KSR) (Life Technologies). Both, naive mESCs and EpiLCs, were cultured on 0.2% gelatin-treated flasks. The media of EpiLCs was changed once after 24 h and all cells were harvested after 48 h. Cells were tested negative for Mycoplasma contamination by PCR.

### Identical culture conditions for mouse formative and primed as well as human ESCs

129S2C1a mouse EpiSCs^62^ and J1 EpiLCs that were compared directly to mEpiSCs were cultured in UPPS medium consisting of StemMACS iPS Brew XF (Miltenyi Biotec) supplemented with 1 µM IWR-1 (Sigma) and 0.5 µM CHIR (Tocris)^86^. Human ESCs H9 were kept under the same cell culture conditions as mouse EpiSCs. ESCs, EpiSCs, and compared EpiLCs were cultured on plates coated with Matrigel (Corning) diluted 1:100 in DMEM/F-12 (Thermo Fisher Scientific).

For all experiments, cells were differentiated/cultured in three independent flasks and are therefore considered to be 3 biological replicates. Cells were split upon harvesting for total proteome (5 x 10^6^ cells per replicate) and chromatome (15 x 10^6^ cells per replicate) analyses and flash-frozen. Cell amounts for both experiments could be reduced by at least a factor of five. The following descriptions are based on the above-mentioned amounts. Systematic downscaling showed that as few as 1 x 10^4^ to 1 x 10^5^ cells per replicate may suffice (see also Methods Details).

### Total proteome sample preparation

Previously flash-frozen samples were quickly placed on ice and pellets were solubilized in 200 µL lysis buffer (6 M guanidinium Chloride, 100 mM Tris–HCl pH 8.5, 2 mM DTT) and heated for 10 min at 99°C under constant shaking at 1,400 rpm. Subsequently, samples were sonicated at 4°C in 30 s on/off intervals for 15 cycles using a Bioruptor ® Plus sonication instrument (Diagenode) at high-intensity settings. If the viscosity of the samples was sufficiently reduced, protein concentrations were estimated, otherwise, sonication was repeated. For concentration measurements, the Pierce™ BCA Protein Assay Kit (23225, Thermo Fisher Scientific) was employed following the manufacturer’s instructions. After at least 20 min of incubation with 40 mM chloroacetamide, 30 µg of each proteome sample was diluted in a 30 µL lysis buffer supplemented with CAA and DTT. Samples were diluted in 270 µL digestion buffer (10% acetonitrile, 25 mM Tris–HCl pH 8.5, 0.6 μg Trypsin/sample (Pierce™ Trypsin Protease, 90058, Thermo Fisher Scientific) and 0.6 μg/sample LysC (Pierce™ LysC Protease, 90051, Thermo Fisher Scientific) and proteins digested for 16 h at 37°C with constant shaking at 1,100 rpm.

To stop protease activity 1% (v/v) trifluoroacetic acid (TFA) was added the next day and samples were loaded on self-made StageTips consisting of three layers of SDB-RPS matrix (Empore)^87^ that were previously equilibrated by 0.1% (v/v) TFA. After loading, two washing steps with 0.1% (v/v) TFA were scheduled and peptides were eluted by 80% acetonitrile and 2% ammonium hydroxide. Upon evaporation of the eluates in a SpeedVac centrifuge, samples were resuspended in 20 µL 0.1% TFA and 2% acetonitrile. After complete solubilization of peptides by constant shaking for 10 min at 2,000 rpm, peptide concentrations were estimated on a Nanodrop^TM^ 2000 spectrophotometer (Thermo Fisher Scientific) at 280 nm.

### Chromatin aggregation capture

Previously flash-frozen samples were quickly placed on ice and pellets were solubilized in 1 mL ChAC lysis buffer (20 mM HEPES pH 7.4, 10 mM NaCl, 3 mM MgCl_2_, 0.1% NP40, freshly added 1× cOmplete™ EDTA-free Protease Inhibitor Cocktail (04693132001, Roche)) and incubated for 10 min on ice. Nuclei were pelleted by centrifugation (2,300 g, 5 min, 4°C) and the supernatant was discarded. In the differential fraction analysis (Fig. 2a), the supernatant was saved as the cytosolic fraction. Upon a second wash of the nuclei pellet with the ChAC lysis buffer, the nuclei were taken into 3 mL crosslinking buffer (PBS pH 7.4 (806552, Sigma), 1x cOmplete™ EDTA-free Protease Inhibitor Cocktail). Formaldehyde (28906, Thermo Fisher Scientific) was added to a final concentration of 1% and samples were incubated for 10 min on an orbital shaker at room temperature. Excess formaldehyde was then quenched by 125 mM Glycine for 5 min and crosslinked cells were washed twice with ice-cold PBS. Nuclei were lysed in 300 µL SDS buffer (50 mM HEPES pH 7.4, 10 mM EDTA pH 8.0, 4% UltraPure™ SDS Solution (24730020, Invitrogen), freshly added 1× cOmplete™ EDTA-free Protease Inhibitor Cocktail) by gentle pipetting. After 10 min incubation at room temperature, 900 µL freshly prepared Urea buffer (10 mM HEPES pH 7.4, 1 mM EDTA pH 8.0, 8 M Urea (U4883, Sigma)) was added. Tubes were carefully inverted 7 times and centrifuged at 20,000 g and room temperature for 30 min. The supernatant was discarded without perturbing the pellet. The pellet was resuspended in 300 µL Sonication buffer (10 mM HEPES pH 7.4, 2mM MgCl_2_, freshly added 1× cOmplete™ EDTA-free Protease Inhibitor Cocktail). Before sonication, two additional wash steps can be scheduled (one SDS and Urea wash and one SDS only wash)^25^, but to our hands, this did not notably improve the chromatin enrichment efficiency. The chromatin samples were sonicated using a Bioruptor ® Plus at 4°C for 15 cycles (30 s on, 60 s off). The protein concentration was estimated by the Pierce™ BCA Protein Assay Kit.

Next, protein aggregation capture (PAC) was performed. Here 1,000 µg of undiluted Sera-Mag^TM^ beads (1 mg, GE24152105050250, Sigma) per 100 µg chromatin solution were washed three times by 70% acetonitrile. 300 µL of the chromatin solution corresponding to 100 µg was added after the last wash to the beads and chromatome-bead mixtures were vortexed. After 10 min incubation on the bench, the samples were again vortexed and rested on the bench. A first wash followed this with 700 µL 100% acetonitrile, a second wash with 1 mL 95% acetonitrile, and a third wash with 1 mL 70% ethanol. The remaining ethanol was allowed to evaporate and beads were resuspended in 400 µL 50 mM HEPES pH 8.5 supplemented with fresh 5 mM TCEP and 5.5 mM CAA. Samples were incubated for 30 min at room temperature upon which LysC (1:200) and Trypsin (1:100) were added. Proteins were digested overnight at 37°C. From this step on, samples were treated exactly like the total proteome samples.

### Chromatin aggregation capture of < 1 million cells

Chromatin aggregation capture for sub-million amounts of cells was performed with some additional modifications to the standard protocol. Here, cells were directly harvested into a DNAse-/RNase-free 1.5 mL tube (0030108051, Eppendorf). Nuclei were then lysed in 0.5 mL of ChAC-lysis buffer and the nuclei pellet was resuspended in 666 µL crosslinking buffer. After crosslinking with 1% formaldehyde and subsequent formaldehyde quenching with 125 mM Glycine, the chromatin extraction was performed again by SDS and Urea washes with careful pipetting so that nothing would stick to the pipette tip. Of note, with < 100,000 cells the chromatin is not visually pelleted but rather a smear that spreads at the wall of the tube. For 10,000 cells even this smear is not visible anymore and it is advised to use a thermal shaker at 1,500 rpm instead of pipetting. For 10,000 to 250,000 cells the protein yield after sonication was between 10-16 µg. Here, we used 10 µg as input for the PAC purification and 1,500 µg magnetic beads per replicate since smaller amounts require a higher bead-to-protein ratio^28^. After the peptide cleanup, these samples were resuspended in 8 µL of 0.1% TFA and 2% acetonitrile.

### Nanoflow LC-MS/MS measurements for proteomes and chromatomes

Peptides were separated prior to MS by liquid chromatography on an Easy-nLC 1200 (Thermo Fisher Scientific) on in-house packed 50 cm columns of ReproSilPur C18-AQ 1.9-µm resin (Dr. Maisch GmbH). By employing a binary buffer system (buffer A: 0.1% formic acid and buffer B: 0.1% formic acid and 80% acetonitrile) with successively increasing buffer B percentage (from 5% in the beginning to 95% at the end) peptides were eluted for 120 min under a constant flow rate of 300 nL/min. Via a nanoelectrospray source, peptides were then injected into an Orbitrap Exploris™ 480 mass spectrometer (Thermo Fisher Scientific). Samples were scheduled in triplicates and a subsequent washing step while the column temperature was constantly at 60°C. Thereby the operational parameters were monitored in real-time by SprayQc.

DDA-based runs consisted of a top12 shotgun proteomics method within a range of 300-1,650 *m*/*z*, a default charge state of 2, and a maximum injection time of 25 ms. The resolution of full scans was set to 60,000 and the normalized AGC target was set to 300%. For MS2 scans the orbitrap resolution was set to 15,000 and the normalized AGC target to 100%. The maximum injection time was 28 ms.

DIA-based runs employed an orbitrap resolution of 120,000 for full scans in a scan range of 350-1,400 *m*/*z*. The maximum injection time was set to 45 ms. For MS2 acquisitions the mass range was set to 361-1,033 with isolation windows of 22.4 *m*/*z*. A window overlap of 1 *m*/*z* was set as default. The orbitrap resolution for MS2 scans was at 30,000, the normalized AGC target at 1,000%, and the maximum injection time at 54 ms. The tested DIA methods varied within the range of the isolation windows which were 37.3 *m*/*z* for in total of 18 windows and 16.8 *m*/*z* for in total of 40 windows.

### MS data quantification

DIA-NN-based analysis of raw MS data acquired in DIA mode was performed by using version 1.7.17 beta 12 in “high accuracy” mode. Instead of a previously measured precursor library, spectra and RTs were predicted by a deep learning-based algorithm and spectral libraries were generated from FASTA files. Cross-run normalization was established in an RT-dependent manner. Missed cleavages were set to 1. N-terminal methionine excision was activated and cysteine carbamidomethylation was set as a fixed modification. Proteins were grouped with the additional command “--relaxed-prot-inf”. Match-between runs was enabled and the precursor FDR was set to 1%.

The DIA raw files were analyzed with the Spectronaut Pulsar X software package (Biognosys, version 14.10.201222.47784)^32^ applying the default Biognosys factory settings for DIA analysis (Q-value cutoff at precursor and protein level was set to 0.01). Imputation of missing values was disabled.

The DDA raw files were analyzed with MaxQuant 1.6.11.0^88^. “Match between runs” was enabled and the FDR was adjusted to 1%, including proteins and peptides. The MaxLFQ algorithm was enabled for the relative quantification of proteins^89^. Contaminants were defined by using the Andromeda search engine^90^.

### Statistical analyses

Downstream analysis of raw data output was performed with Perseus (version 1.6.0.9)^91^. For the calculation of CVs, proteins or precursors with less than 2 out of 3 valid values were filtered out. For GO term counts the filtering was more strict and 3 out of 3 valid values were required. GO enrichment analyses of differentially enriched proteins (Fig. 2a) were performed against the background of total identified proteins by employing a Benjamini-Hochberg FDR-corrected Fisher’s Exact test. The analysis was thereby performed individually for each cluster. The functional enrichment analysis of proteins enriched by ChAC-DIA vs total proteome was performed by ranking proteins according to their enrichment in the ChAC-DIA fraction. The functional enrichment analysis was thereby based on STRING^92^.

Student’s t-tests were performed after imputation of missing values. The latter was always performed based on a Gaussian distribution relative to the standard deviations of measured values (width of 0.2 and a downshift of 1.8 standard deviations). Both, one- and two-sided t-tests were calculated with a permutation-based FDR of 0.05 and an s0 = 1 if not otherwise declared. For the multiple sample test based on an ANOVA (Fig. 2a) we chose a minimal 1.5-fold change but did not apply imputation. Student’s t-tests of normalized chromatomes were performed after calculating pairwise differences of ChAC-DIA and total proteome values. The complete catalog of proteins found in the naive, formative, and primed states can be found in Supplementary Table 3.

Correlations between samples in the differential fraction analysis experiment were calculated with Perseus, and the correlations between transcriptomes, proteomes, and chromatomes were calculated with GraphPad Prism (version 9.1.0).

### Web application development

Row-normalized z-scores for each significantly changing protein across the ChAC-DIA purification steps were generated for an interactive profile plot representation of the data. Significant chromatome and chromatin affinity changes during pluripotency were represented in an interactive heatmap as mean row differences of Log2 intensities and mean row differences of z-scores, respectively. The high confidence chromatome comparisons between hESCs and mouse PSCs are based on mean row Log2 intensities.

The web application was programmed using R Shiny with the following libraries besides base R packages for data processing and visualization: shiny (1.7.1), shinydashboard (0.7.2), shinyHeatmaply (0.2.0), plotly (4.10.0), heatmaply (1.3.0) and png (0.1-7). From the tidyverse (1.3.1) family we further utilized tidyr (1.2.0), dplyr (1.0.9), and ggplot2 (3.3.6).

### Data availability

The mass spectrometry proteomics data have been deposited to the ProteomeXchange Consortium via the PRIDE^85^ partner repository with the dataset identifier PXD034448. To access the data, please use the reviewer account with the following login details:

Username:

Password:

To make the files better comprehensible, they have been assigned to experiments (Raw data list, see PXD034448). Source data are provided in this paper. The used RNA-Seq dataset is derived from the ArrayExpress with the following accession code: E-MTAB-6797.

### Code availability

The underlying custom code for the provided web application, accessible on https://ugur-enes.shinyapps.io/Chromatome_Atlas/.

## Acknowledgments

We thank Dr Igor Paron for excellent MS technical assistance, Dr Johannes Bruno Müller-Reif for columns, and Jeannette Koch and Ruzica Barisic for technical assistance. We thank Dr Christopher B. Mulholland, Dr Michael D. Bartoschek as well as Dr Carsten Marr for valuable and insightful scientific discussions. We thank Dr Vadim Demichev for generously providing access to the most recent version of DIA-NN and for answering questions on setting up DIA-NN. We thank Dr Ludovic Vallier and Dr Derk ten Berge for kindly providing mEpiSCs. E.U. gratefully acknowledges the International Max Planck Research School for Molecular Life Sciences (IMPRS-LS) and the research training group 1721 (RTG 1721) for training and support. This work was funded by the Deutsche Forschungsgemeinschaft (DFG, German Research Foundation) – Project-ID 213249687 – SFB1064 to H.L. (A17) and S.B. (A22).

## Author information

These authors contributed equally: Alexandra de la Porte and Sebastian Bultmann.

These authors jointly supervised: Matthias Mann, Michael Wierer and Heinrich Leonhardt.

## Contributions

E.U. and M.W. designed the study; H.L., M.W., and M.M. supervised the study. E.U. performed all experiments and MS data analysis, and E.U., H.L., and M.W. interpreted the data. E.U., H.L., M.W., and S.B. wrote the manuscript and prepared the figures. A.d.l.P. established the same culture conditions for human and mouse ESCs and conducted the cell culture under the supervision of M.D. E.U. and S.B. programmed the web application. All authors discussed the results, then read and approved the manuscript.

## Ethics declarations

The authors declare no competing interests.

## Supplementary information

Supplementary Fig. 1–10

Supplementary Table 1. Benchmarking data of Chromatin Aggregation Capture (ChAC) followed by data-independent MS acquisition (DIA), Related to Fig. 1 b-f, Supplementary Fig. 1 a-h and 2 a-d

Supplementary Table 2. Chromatome map of naive mESCs based on ChAC-DIA, Related to Fig. 2 a-e and Supplementary Fig. 3 a-f

Supplementary Table 3. Chromatome atlas of mouse pluripotency phases, Related to Fig. 3 b-e, 4 e,f and Supplementary Fig. 4-8

Supplementary Table 4. Comparison of human and mouse pluripotency, Related to Fig. 5 a-h and Supplementary Fig. 10 a-h

## Notes

https://ugur-enes.shinyapps.io/Chromatome_Atlas/

### Competing Interest Statement

The authors have declared no competing interest.

