## Supplementary information for "Comprehensive chromatin proteomics resolves functional phases of pluripotency"

**This PDF includes:**

Supplementary Fig. 1 (Related to Fig. 1)
Supplementary Fig. 2 (Related to Fig. 1)
Supplementary Fig. 3 (Related to Fig. 2)
Supplementary Fig. 4 (Related to Fig. 3)
Supplementary Fig. 5 (Related to Fig. 3)
Supplementary Fig. 6 (Related to Fig. 3)
Supplementary Fig. 7 (Related to Fig. 3)
Supplementary Fig. 8 (Related to Fig. 3)
Supplementary Fig. 9 (Related to Fig. 3)
Supplementary Fig. 10 (Related to Fig. 5)

#### Supplementary Fig. 1 (Related to Fig. 1)

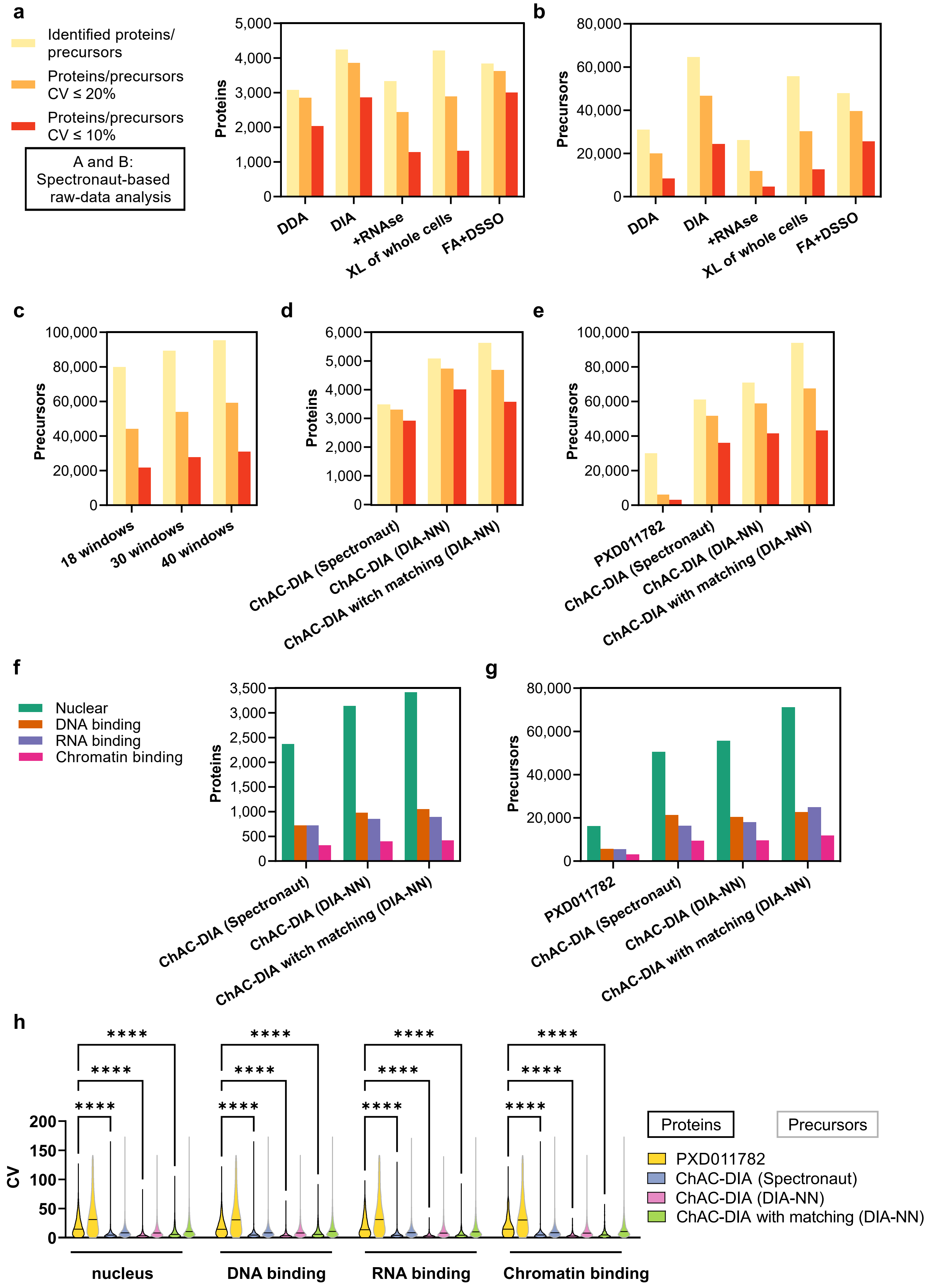

#### Supplementary Fig. 2 (Related to Fig. 1)

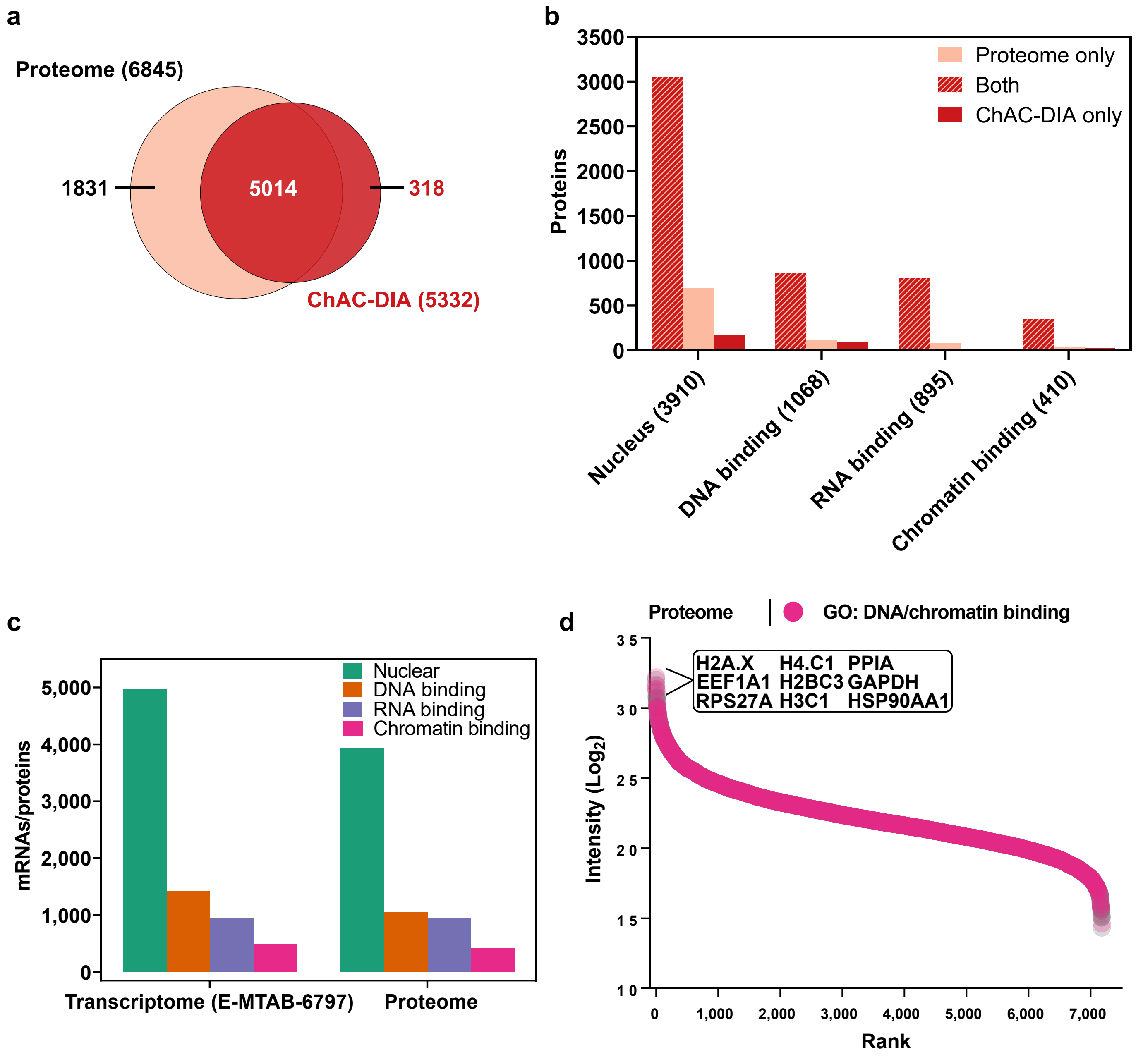

### Supplementary Fig. 3 (Related to Fig. 2)

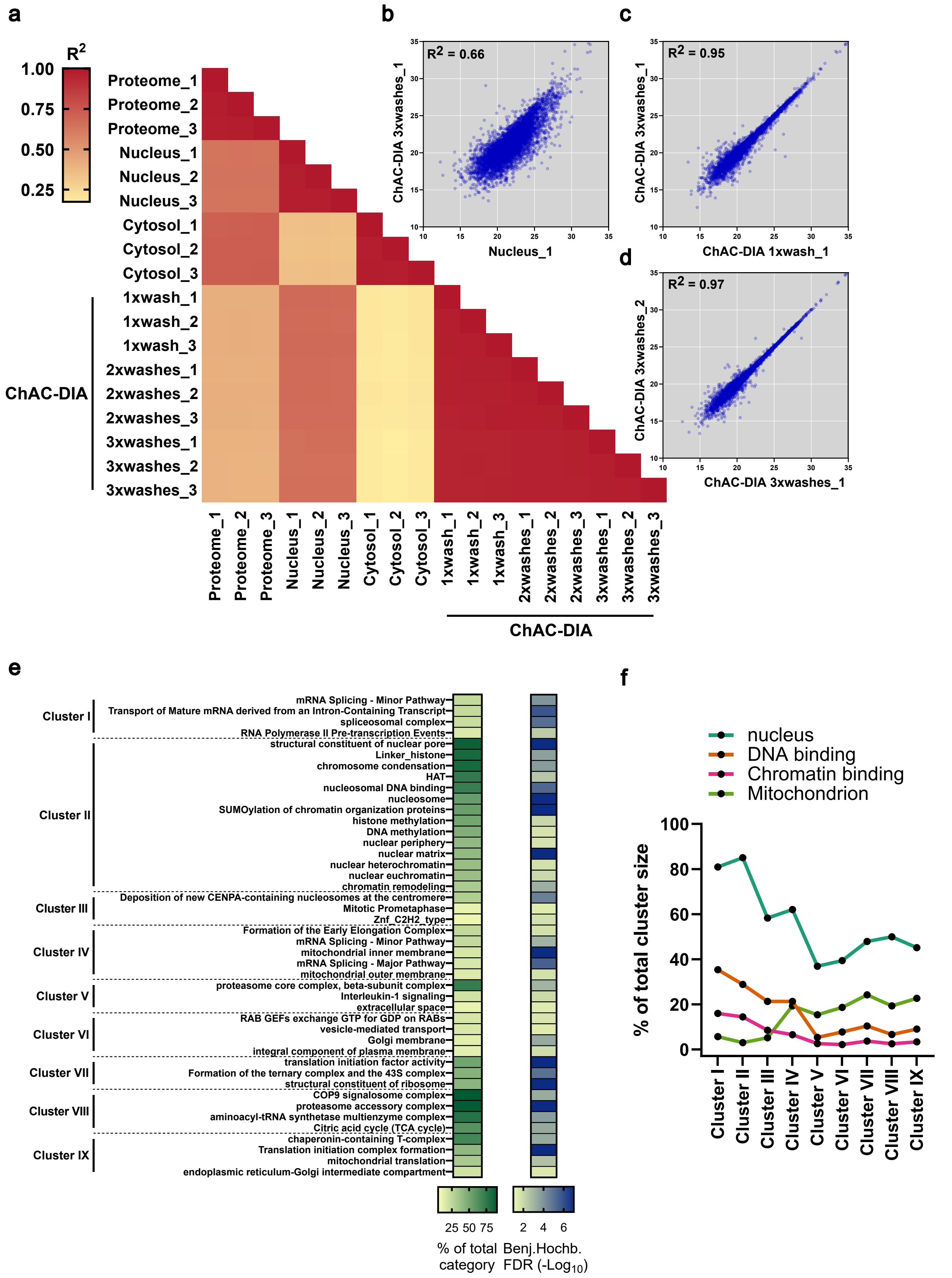

**Supplementary Fig. 4 (Related to Fig. 3)**

### Pluripotency/differentiation marker proteins

a) ANOVA significant (FDR < 0.05, fold change difference ≥ 2)

b) Absent in at least one pluripotency phase

c) Not ANOVA significant between any pluripotency phase

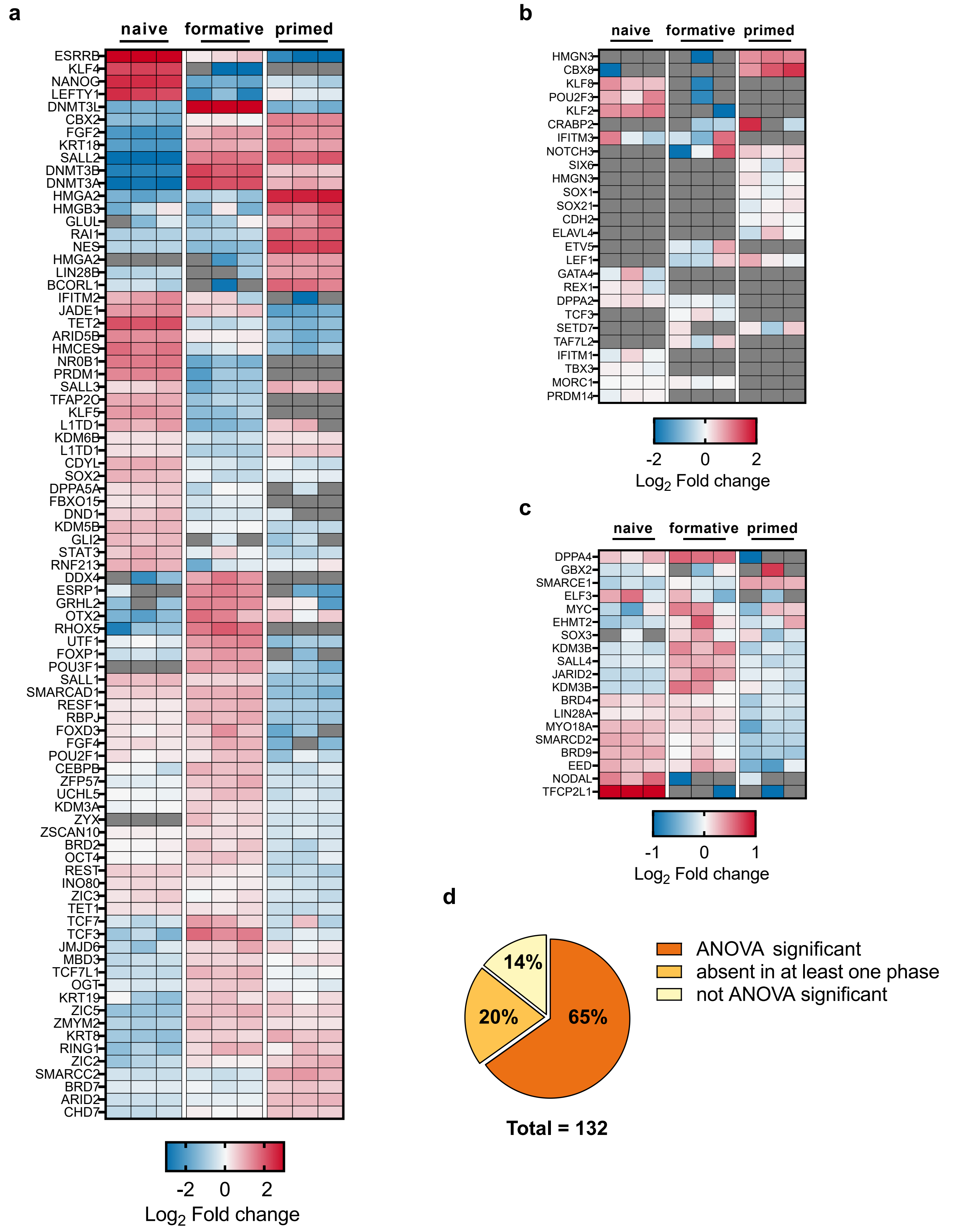

**Supplementary Fig. 5 (Related to Fig. 3)**

### GO "Transcription factor activity"

a) **ANOVA significant (FDR < 0.05, fold change difference ≥ 2)**

b) **Absent in at least one pluripotency phase**

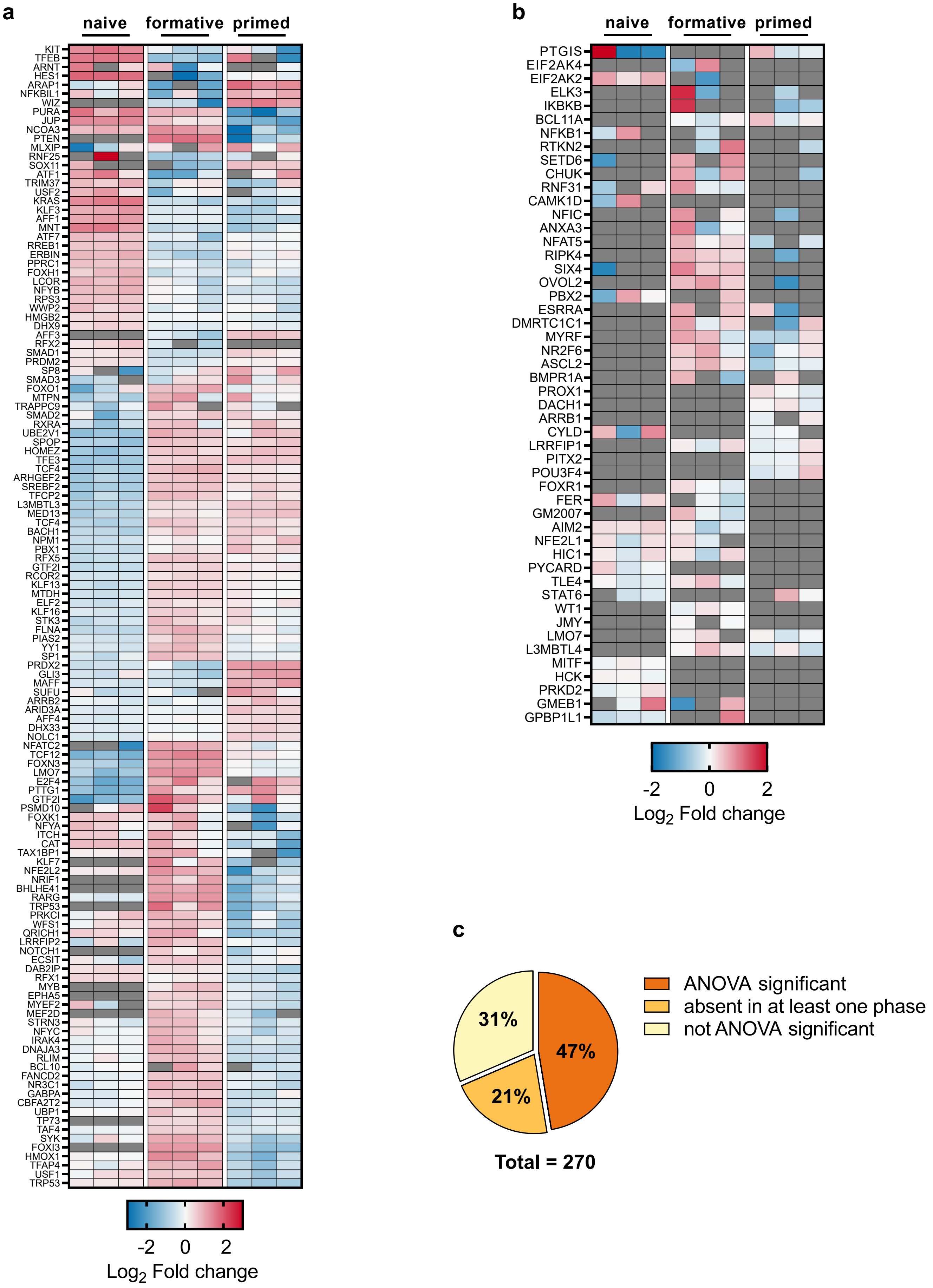

Supplementary Fig. 6 (Related to Fig. 3)

### Proteins related in epigenetic regulation

a) ANOVA significant (FDR < 0.05, fold change difference ≥ 2)

b) Absent in at least one pluripotency phase

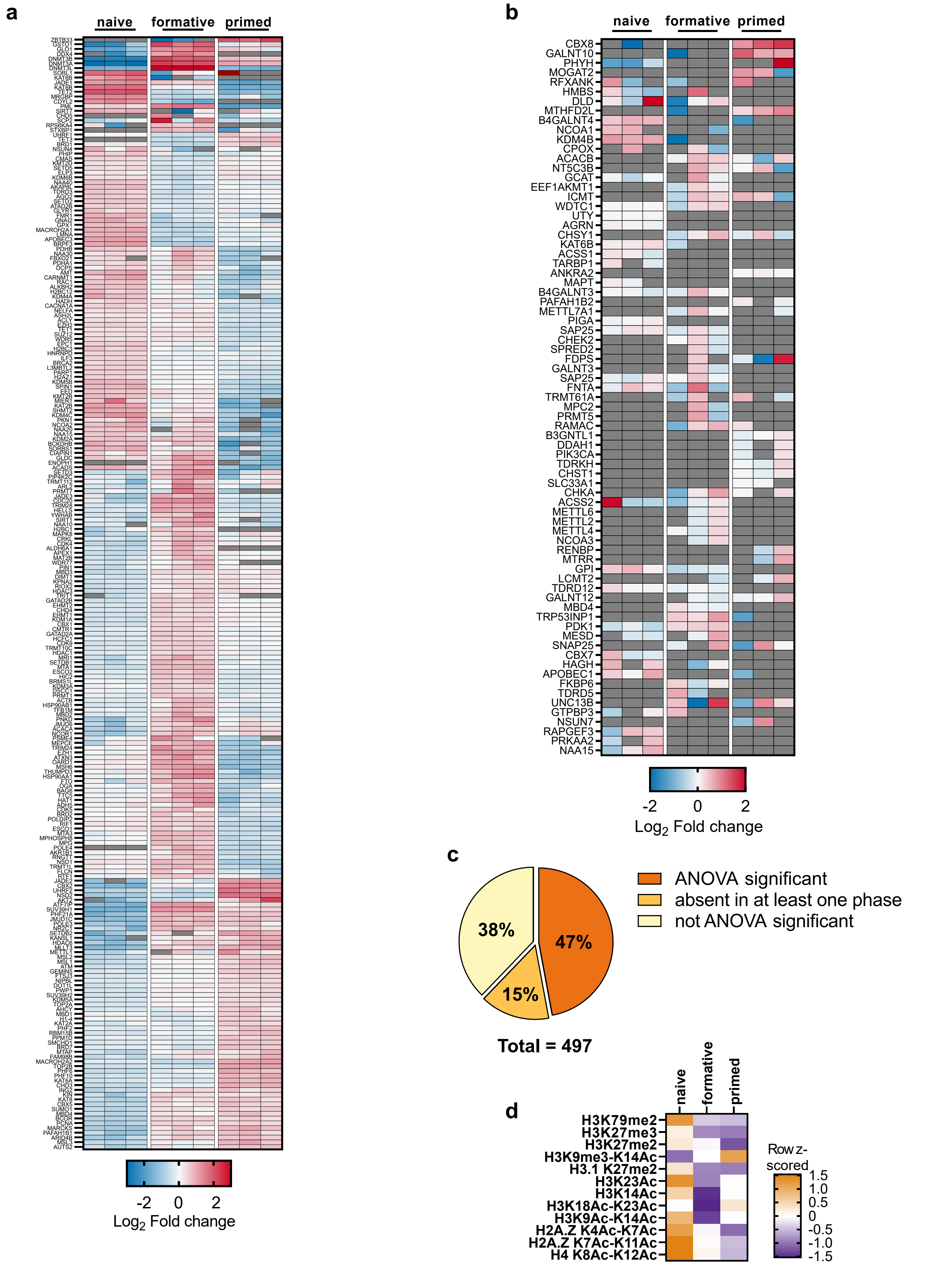

**Supplementary Fig. (Related to Fig. 3)**

### Zinc finger (domain containing) proteins

**a) ANOVA significant (FDR < 0.05, fold change difference ≥ 2)**

**b) Absent in at least one pluripotency phase**

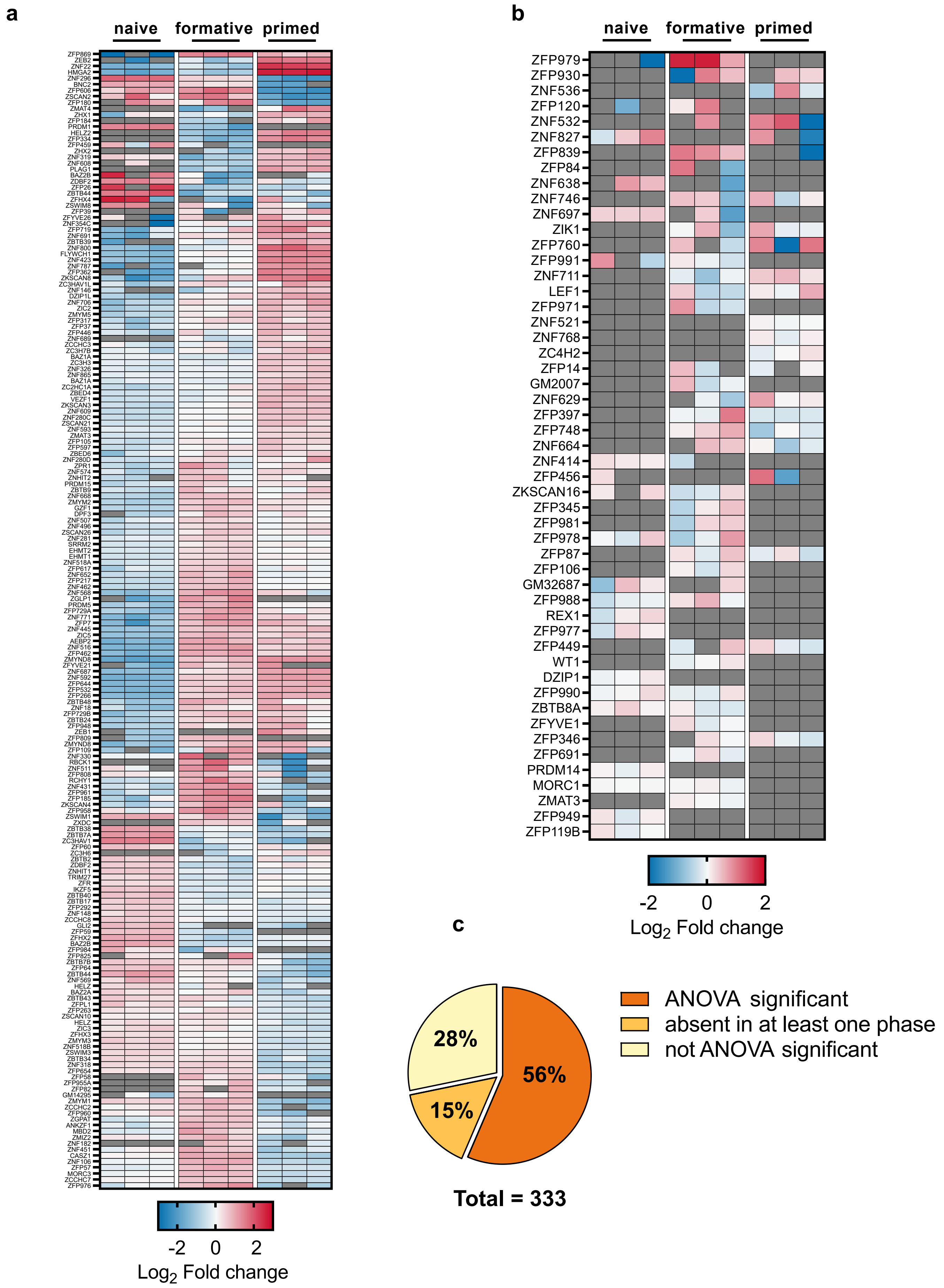

**Supplementary Fig. 8 (Related to Fig. 3)**

### Chromatin remodeler

**a) ANOVA significant (FDR< 0.05, fold change difference ≥ 2)**

**b) Absent in at least one pluripotency phase**

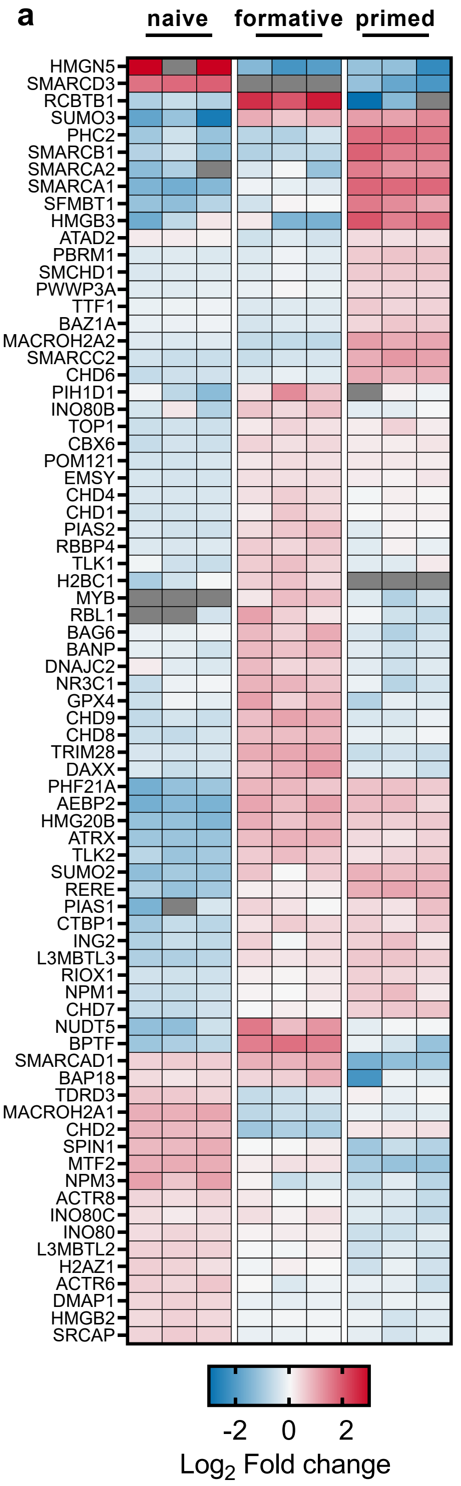


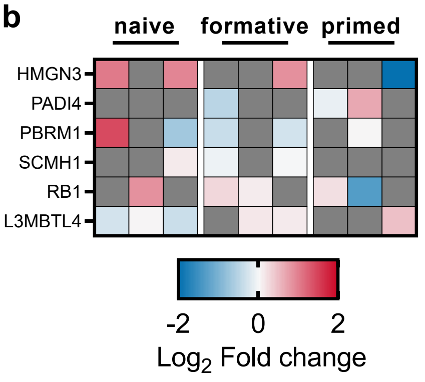


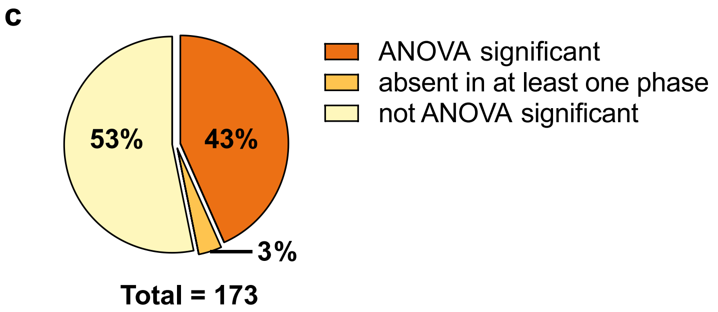

**Supplementary Fig. 9 (Related to Fig. 3)**

### Chromatin binding complexes

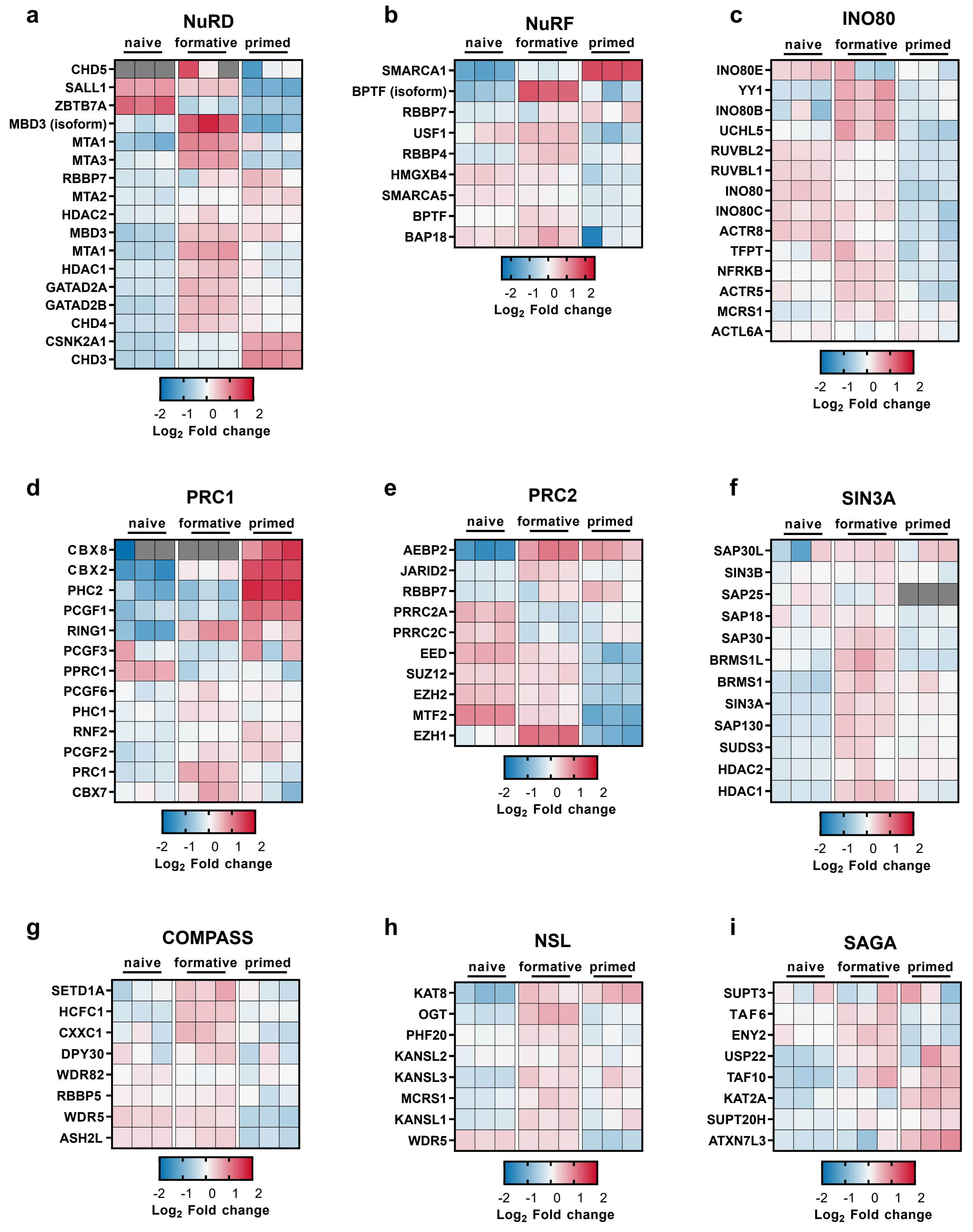

**Supplementary Fig. 10 (Related to Fig. 5)**

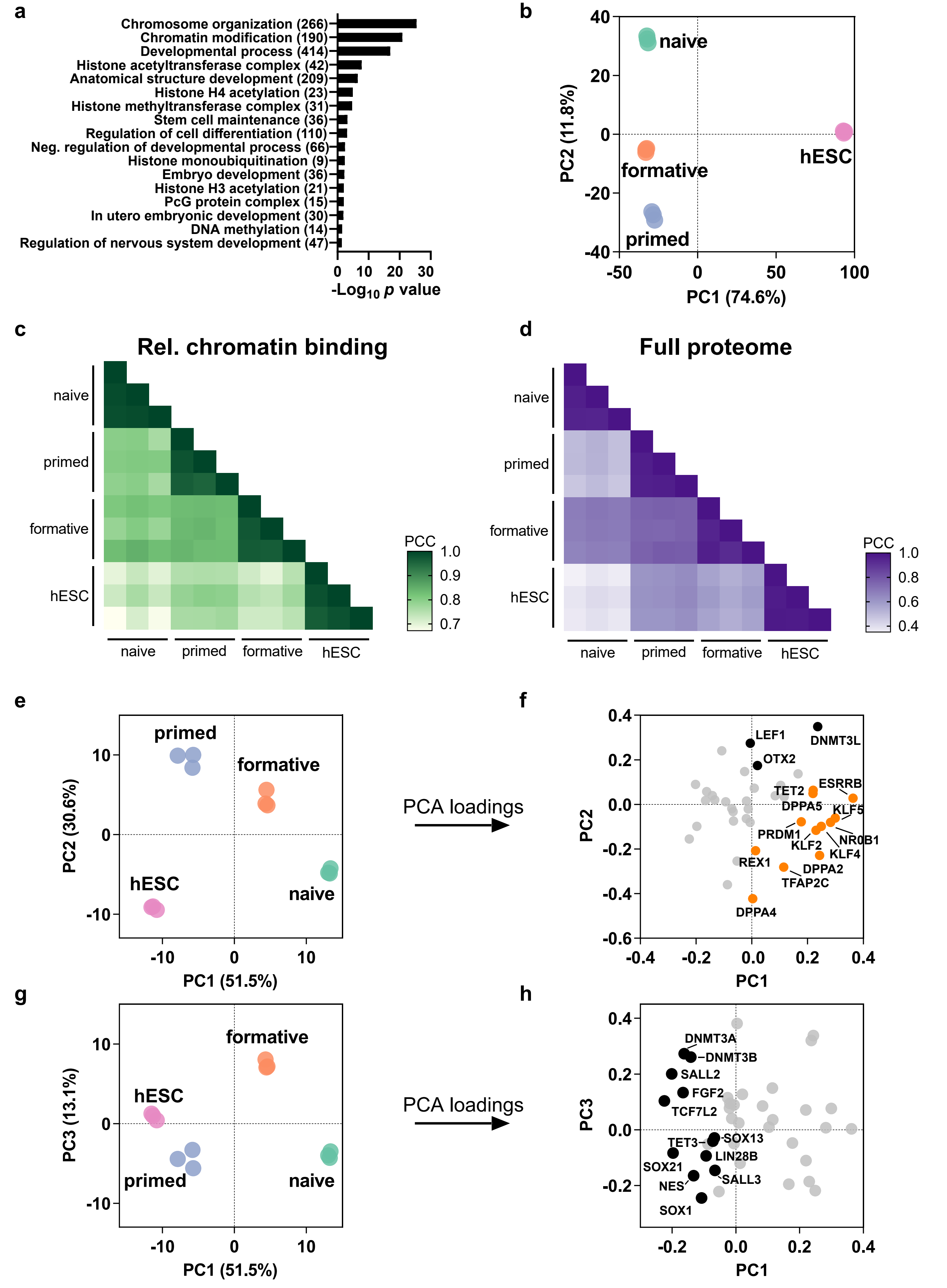

#### Supplemental Figure legends

##### Supplementary Fig. 1. Evaluation of ChAC-DIA improvements, Related to Fig. 1.

**a, b** Total numbers of identified proteins (**a**) or precursors (**b**) with representation of the percentage coefficient of variation (CV) below 20% and 10%. We performed our protocol with changing only one parameter at once. DDA: same ChAC workflow, but without DIA. DIA: PAC step is omitted (instead, acetone precipitation is performed). +RNase: additional incubation of nuclei with RNaseA for 15 min at 37°C. XL of whole cells: FA crosslinking is performed before nuclei isolation. FA+DSSO: double crosslinking with FA and DSSO. The data was analyzed with Spectronaut. Experimental conditions were kept comparable by using the same cell pool. Briefly, DIA improved the protein identification rate by 37.9% with constant CVs around 7.3% compared to just DDA. Despite more than doubling precursor numbers, CVs on peptide-level were even reduced from 15.6% for DDA to 12.7% for DIA. Strikingly, CVs were further improved by the additional PAC step (median CVs for proteins 4.1% and for precursors 9.6%). Moreover, additional RNAse addition prior to nuclei lysis and the formaldehyde crosslinking of whole cells instead of nuclei impaired CVs especially on precursor level (22.2% and 18.4%, respectively). A combination of DIA (see **d** for optimizations) and PAC after crosslinking of purified nuclei without RNaseA addition therefore gave the best results in terms of sensitivity and reproducibility. **c** Effect of precursor isolation window numbers in DIA on total precursor identifications and CV. The data was analyzed with DIA-NN. MS2 resolution was constant at 30,000. Here, we found that 30 or 40 isolation windows outperform 15 windows regarding total precursor identifications by 11.8% and 19.3%, respectively, while keeping the CVs constant. **d, e** Total numbers of identified proteins (**d**) or precursors (**e**) with representation of the percentage CVs below 20% and 10% (corresponding experiment to **Fig. 1b**). We compare a previous study (PRIDE: PXD011782) to ChAC-DIA quantified in directDIA mode by either Spectronaut or DIA-NN with or without matching across different purification fractions mentioned in **Fig. 2a**. **f, g** Total numbers of proteins (**f**) or precursors (**g**) falling into a gene ontology (GO) category (corresponding experiment to **Fig. 1c**). **h** Distribution of CVs of proteins selected by GO category. Asterisks represent digits after the decimal point of 0.05. For better readability only comparisons on protein-level are shown. However, each comparison between PXD011782 and any given ChAC-DIA analysis method was to the same extent significant on precursor- level.

##### Supplementary Fig. 2. Comparison of chromatome to proteome and transcriptome, Related to Fig. 2.

**a** Venn diagram of proteins identified in proteome and chromatome of naive PSCs. Biological replicates: n = 3. **b** Numbers of proteins falling into a GO category and that are either identified in the proteome and chromatome or in one of the experiments exclusively. **c** Numbers of proteins falling into a GO category in the proteome or transcriptome of naïve PSCs. **d** Protein abundance rank based on the naive PSC proteome. Chromatin binding and DNA-binding proteins are highlighted in magenta. Displayed proteins indicate highest ranked 9 proteins.

**Supplementary Fig. 3. Reproducibility of differential fraction analysis during ChAC- DIA, Related to Fig. 2.**

**a-d** Unsuprevised hierarchical clustering of $R^2$ values (**a**) and scatter plots of nucleus vs ChAC-DIA (3x washes) (**b**), ChAC-DIA 1x wash vs 3x washes (**c**) and two replicates of ChAC-DIA (3x washes) (**d**). **e** Fisher's exact test to assess enriched GO terms of significantly enriched proteins in each cluster based on unsupervised hierarchical clustering in **Fig. 2a**. P values are colour coded ($-Log_{10}$) and dot diameters correspond to group sizes ($Log_2$). **f** Percentage of a given GO category from the total cluster size of each cluster.

**Supplementary Fig. 4. Complete chromatome list of pluripotency or differentiation associated proteins, Related to Fig. 3.**

**a-c** Heatmap representation of $Log_2$ FCs for significant differences between pluripotency phases (**a**), proteins absent in at least one pluripotency phase (**b**) or no ANOVA-based significant differences (**c**). **d** Total group size and percentage of (non-)significant differences.

**Supplementary Fig. 5. Complete chromatome list of proteins annotated with „Transcription factor activity", Related to Fig. 3.**

**a, b** Heatmap representation of $Log_2$ FCs for significant differences between pluripotency phases (**a**) or proteins absent in at least one pluripotency phase (**b**). **c** Total group size and percentage of (non-)significant differences.

**Supplementary Fig. 6. Complete chromatome list of proteins associated with epigenetic regulation, Related to Fig. 3.**

**a, b** Heatmap representation of $Log_2$ FCs for significant differences between pluripotency phases (**a**) or proteins absent in at least one pluripotency phase (**b**). **c** Total group size and percentage of (non-)significant differences. **d** Selection of identified and ANOVA-significant histone posttranslational modifications (PTMs) identified by ChAC-DIA. Data was analyzed by Spectronaut, column-wise normalized to median column intensity and subsequently row z-scored and averaged across triplicates.

**Supplementary Fig. 7. Complete chromatome list harboring a Zinc finger domain, Related to Fig. 3.**

**a, b** Heatmap representation of $Log_2$ FCs for significant differences between pluripotency phases (**a**) or proteins absent in at least one pluripotency phase (**b**). **c** Total group size and percentage of (non-)significant differences.

**Supplementary Fig. 8. Complete chromatome list of proteins annotated with „chromatin remodeler" or „chromatin organization", Related to Fig. 3.**

**a, b** Heatmap representation of $Log_2$ FCs for significant differences between pluripotency phases (**a**) or proteins absent in at least one pluripotency phase (**b**). **c** Total group size and percentage of (non-)significant differences.

**Supplementary Fig. 9. Examples of chromatin-associated complexes, Related to Fig. 3.**

**a-i** Heatmap representation of $Log_2$ FCs for changes between pluripotency phases.

**Supplementarey Fig. 10. Comparison of proteomes and chromatomes between mouse naive, formative and primed PSCs as well as hESCs, Related to Fig. 5.**

**a** Fisher's exact results obtained from comparing the shared high confidence chromatome across all four tested cell lines against the total set of high confidence chromatome binders. Numbers in brackets represent groups size. **b** PCA of the high confidence chromatomes of the tested three mouse cell lines. **c, d** Pearson correlations of relative chromatin binding (**c**) or full proteomes (**d**) filtered for pluripotency or differentiation markers as in **Fig. 5d**. Underlying data was filtered for only valid values (**c**) or at least 6 valid values in total and missing values were imputed based on a gaussian distribution relative to the standard deviations of measured values (width of 0.2 and a downshift of 1.8 standard deviations) (**d**). **e-h** PCA representations of projections (**e, g**) and individual loadings (**f, h**) based on chromatome values of bona fide pluripotency and differentiation markers as represented in **Fig. 5h** and from each mouse PSC and hESCs. **e, g** are based on PC1 and PC2 whereas **f, h** on PC1 and PC3. Orange dots represent pre- implantation markers and black dots post- implantation markers.
